# Detecting altered hepatic lipid oxidation by MRI in an animal model of NAFLD

**DOI:** 10.1101/2023.08.16.553555

**Authors:** Marc McLeod, Mukundan Ragavan, Mario Chang, Rohit Mahar, Anthony Giacalone, Anna Rushin, Max Glanz, Vinay Malut, Dalton Graham, Nishanth E. Sunny, Matthew E. Merritt

## Abstract

Nonalcoholic fatty liver disease (NAFLD) prevalence is increasing annually and affects over a third of U.S. adults. NAFLD can progress to nonalcoholic steatohepatitis (NASH), characterized by severe inflammation and fibrosis. NASH is predicted to become the primary cause of liver transplant by 2030. Although the etiology of NAFLD/NASH is incompletely understood, dysregulated fatty acid oxidation is implicated in disease pathogenesis. Here, we developed a method for estimating hepatic β-oxidation from the metabolism of [D_15_]octanoate to deuterated water and detection with deuterium magnetic resonance methods. Application of this method to perfused liver from a mouse model of NAFLD revealed dysregulated hepatic β-oxidation, findings that we confirmed with *in vivo* imaging. The high-fat diet–induced NAFLD mouse studies indicated that decreased β-oxidative efficiency in the fatty liver could serve as a prognostic indicator of NAFLD progression. Furthermore, our method provides a clinically translatable imaging approach for determining hepatic β-oxidation efficiency.

## Main

The incidence of nonalcoholic fatty liver disease (NAFLD) has risen rapidly over the last few decades and affects more than 30% of adults in the U.S. [1–2]. Of these individuals, ∼20% are expected to progress from nonalcoholic fatty liver (NAFL), defined as 5% or more hepatic fat by mass, to an inflammatory state known as nonalcoholic steatohepatitis (NASH) [1]. Although NAFLD can lead to NASH in some cases, factors triggering the transition are not well established [3]. Unfortunately, ∼17% of individuals with NASH develop debilitating cirrhosis of the liver, and NASH now stands as the #2 cause of liver transplants in the U.S. [1–4]. As viral hepatitis and chronic viral liver disease continue to decline, NASH is predicted to be the number one cause of transplantation by 2030 [4]. A high percentage of obese individuals have fatty liver (70-90%) [1-2, 3, 5-7], and most individuals who develop NASH are obese (81%) [8].

NAFLD diagnoses are based on: (i) pre-screening by body mass index (BMI), along with the presence of insulin resistance and elevated liver enzymes in circulation, (ii) magnetic resonance imaging (MRI) measures of hepatic fat by mass and, (iii) liver stiffness to identify fibrosis [9]. Additionally, liver biopsy is recommended for NAFLD patients to identify inflammation and definitively diagnose NASH [9]. However, the diagnostic accuracy of liver biopsy is limited by sampling error and such biopsies have a known risk of morbidity [10]. Despite this clear protocol for identifying NAFLD and NASH, to date there are no FDA-approved drugs to treat NAFLD progression [5]. Currently, the only clear and effective guidance is weight loss and lifestyle adjustment, a guideline that 80–90% of obese patients fail to achieve and maintain over a 10-year period [1-3, 11]. Although more than 50 compounds have been tested in clinical trials, only two have published positive data in stage 3 clinical trials, necessitating in-depth mechanistic research [12–14].

The accumulation of excess hepatic lipids is associated with the onset of inflammation [1-3, 9, 5-7]. However, the metabolic mechanisms underlying inflammatory onset are still unclear. Lipid accumulation, especially of free fatty acids, leads to endoplasmic reticulum (ER) stress and lipoapoptosis that result from increased ceramide production [15–16]. In some models of NAFLD, decreased oxidative metabolism has been used to infer reduced fatty acid β-oxidation as a culprit for the accumulation of toxic levels of lipids [17]. However, in several obesity models of NAFLD, higher hepatic lipids correlate with increased oxidative metabolism [18–21] potentially to prevent ER stress and lipoapoptosis. However, the increase in oxidative metabolism drives reactive oxygen species (ROS) production that can induce mitochondrial DNA (mtDNA) damage, cause aberrant protein production, and trigger the expression of inflammatory genes [22]. Furthermore, β-oxidation is associated with much greater levels of ROS production than those produced by TCA cycle activity and appears a likely culprit for driving ROS-induced damage during NAFLD progression when lipids are abundant [22]. Despite the importance of hepatic β-oxidation, none of the current studies directly measured hepatic β-oxidation. Instead, these studies inferred β-oxidative flux using fatty acid oxidation products such as ketones or by difference between estimated TCA cycle turnover and oxidation of substrates other than fatty acids. Direct assessment of hepatic β-oxidation would clarify its role in disease pathology.

Deuterated fatty acids produce deuterated water (HDO) as a direct byproduct of β-oxidation, enabling specific, localized spectroscopy or magnetic resonance imaging (MRI) of hepatic β-oxidation [23]. Previously, we used simple MRI protocols to monitor HDO production resulting from glucose metabolism in the rodent brain following bolus administration of [^2^H_7_]glucose [24]. Here, we used [^2^H_15_]octanoate (also referred to as [D_15_]octanoate) as a metabolic contrast agent to study hepatic fatty acid metabolism in a mouse model of high fat diet (HFD)-induced NAFLD [25]. We selected octanoate because it does not require albumin for administration and it readily diffuses into the mitochondria, unlike long chain fatty acids, which are poorly soluble and must be complexed with albumin for administration and require carnitine palmitoyl transferase-1 (CPT-1) to enter the mitochondria [26–27]. Additionally, the rapid uptake of octanoate suppresses activity of the tricarboxylic acid (TCA) cycle [28], enabling selective detection of HDO produced from β-oxidation by deuterium magnetic resonance (DMR). Our method operates in analogy to the glucose tolerance test in which the excess substrate enforces changes in metabolism. We hypothesize this deuterated “lipid challenge” test will be diagnostic of disease state.

## Results

### Deuterium metabolic spectroscopy of [D_15_]octanoate-perfused liver reveals reduced hepatic β-oxidation per g liver protein in a mouse model of NAFLD

As a model, we used control mice on a low-fat diet (LFD, composed of 10% fat by calories) and mice on a HFD (60% fat by calories). The mice on the HFD almost doubled in body weight at the 16-week timepoint (51 +/-2.3 g SD) relative to mice on the LFD (28.6 +/-1.2 g SD) (Student’s T.test, p-value = 1.2*E-6) (Fig. 1A). Additionally, perfused livers from fasted, HFD mice were significantly larger (3.1 g ± 0.25 g, p-value = 4.5*E-5) than their fasted, LFD counterparts (1.6 g ± 0.32 g) and had significantly more total liver protein (Fig. 1B, C), conditions also observed for patients with NAFLD [5].

**Figure 1.**
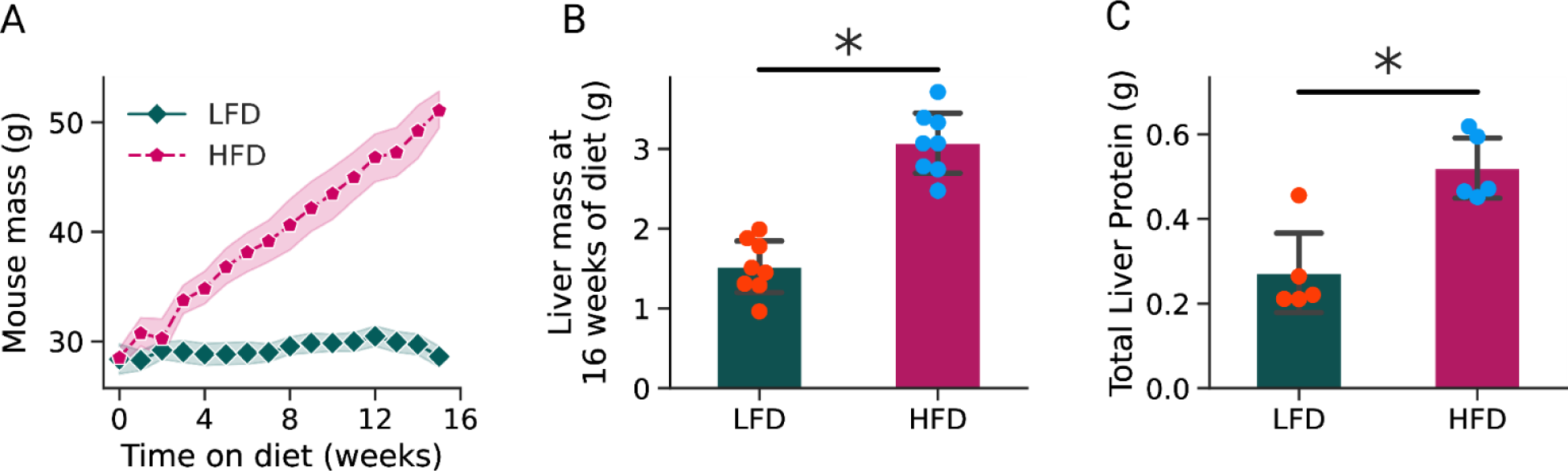
Changes in mouse body weight, liver weight and total liver protein over 16 weeks of HFD or LFD. (A) Weekly mouse mass of 10-week-old C57BL/6J mice after switching to a high fat (60% fat by Kcal) or low fat (10% fat by Kcal) diet. Data are presented as average ± 95% confidence interval (shaded regions). (B, C) Livers of mice on the indicated diets for 16 weeks were collected after *ex vivo* perfusion. Total liver mass was measured (B) and total liver protein was quantified by Bradford assay (C). Error bars represent the mean ± standard deviation. Statistical significance was determined by analysis of covariance for mouse weight gain and Student’s t-test for liver mass and total liver protein (*, p < 0.05

To assess the rate of β-oxidation in the *ex vivo* perfused liver, we measured HDO elution as a byproduct of [D_15_]octanoate oxidation (Fig. 2A, Fig. S1). Because HDO is present at natural abundance (∼.031%) in body water [29], we first assessed HDO by nuclear magnetic resonance (NMR) analysis prior to perfusion of [D_15_]octanoate (Fig. 2B, red trace). We then monitored the increase in HDO after [D_15_]octanoate perfusion (Fig. 2B blue trace), which we set as the start of the period for monitoring HDO. We initially detected an exponential increase in HDO followed by a slower stable increase in HDO that was similar for livers from HFD or LFD mice (Fig. 2C). After normalizing for differences in endogenous (pre-perfusion) HDO, we observed that [D_15_]octanoate accumulated to a greater extent in the livers of the HFD mice than the livers of the LFD mice (Fig. 2D). Normalizing HDO produced by total liver protein showed that the livers of the HFD mice produced less HDO than those of the LFD mice (Fig. 2E). Finally, we normalized the HDO signal to the [D_15_]octanoate signal to assess the balance between fatty acid oxidation and storage during perfusion: Liver that oxidizes more fatty acid than it stores will have a higher ratio. Livers of LFD mice had higher ratios than livers of HFD mice (Fig. 2F). Collectively, these data indicated that livers of HFD mice have reduced efficiency in β-oxidation compared to livers of LFD mice.

**Figure 2.**
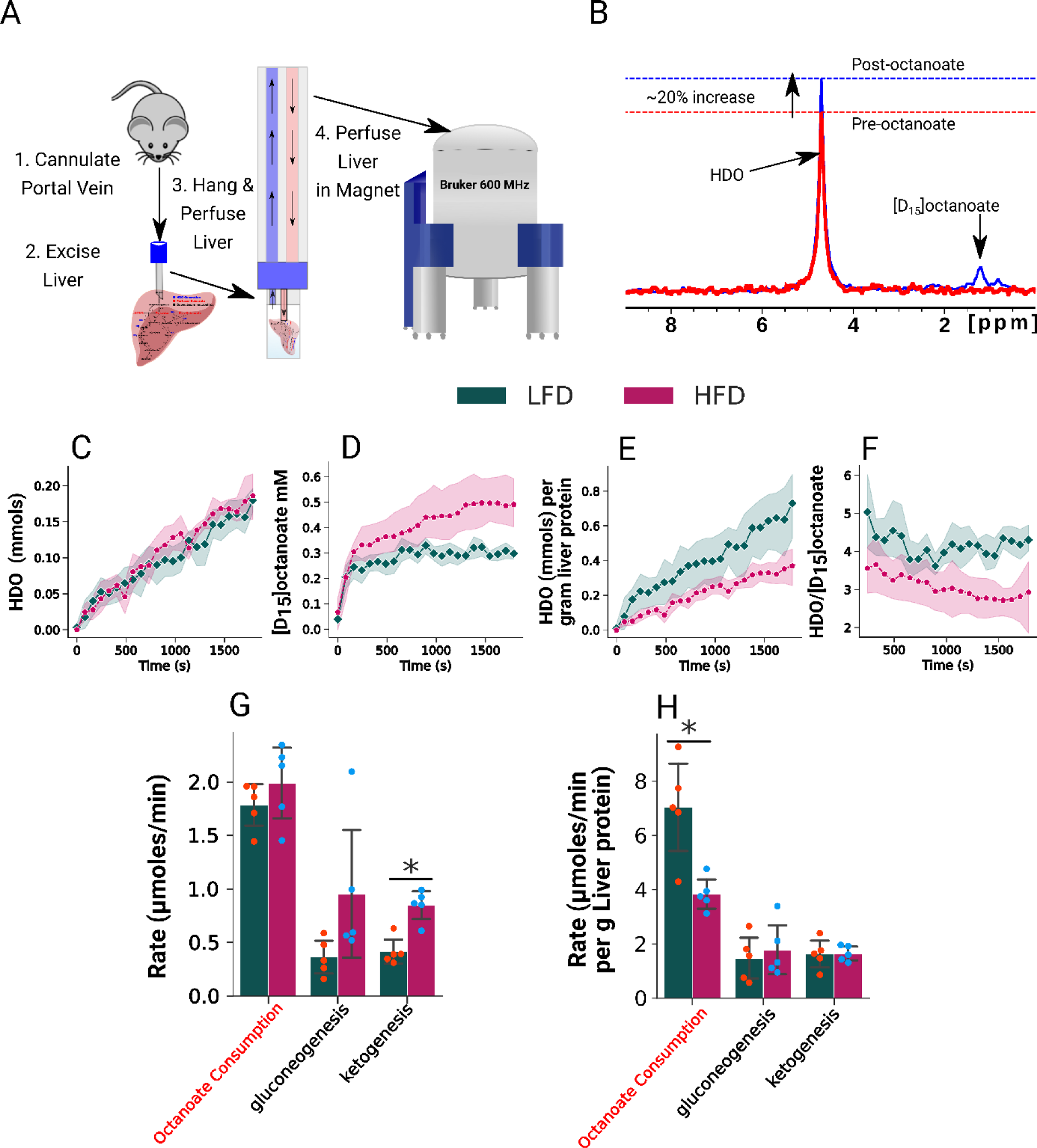
Measures of oxidative metabolism in an *ex vivo* perfused liver system. (A) Diagram of in-magnet perfused mouse liver setup. (B) Representative trace of deuterium magnetic resonance spectroscopy of perfused mouse liver before (red) and 30 minutes after (blue) perfusion with [D_15_]octanoate. (C-F) Quantification of HDO and [D_15_]octanoate peaks presented as average +/-95% confidence interval of individual liver signal in shaded regions. (C) HDO (mmols) produced throughout liver perfusion from [D_15_]octanoate. (D) mM [D_15_]octanoate in liver during perfusion. (E) Total HDO per gram liver protein produced throughout liver perfusion from [D_15_]octanoate. (F) Ratio of HDO signal intensity to [D_15_]octanoate signal intensity. (G) Total and (H) normalized rates of the indicated metabolic processes were determined by GC-MS and quantified from changes in the concentrations of [D_15_]octanoate, glucose, and ketones. Statistical significance was determined by analysis of covariance for NMR quantification and Student’s t-test for GC-MS rate comparisons (*, p < 0.05).

### Analysis of perfusate confirms perturbed β-oxidation in a mouse model of NAFLD

To assess the reliability of our NMR-based β-oxidation analysis, we measured hepatic consumption of [D_15_]octanoate from the efferent perfusate over the course of perfusion by gas chromatography mass spectrometry (GC-MS). Although total [D_15_]octanoate consumption was similar in the livers from the two groups (Fig. 2G), [D_15_]octanoate consumption was significantly lower in the livers from the HFD mice when normalized to total liver protein: HFD mouse livers consumed 3.84 ± 0.60 µmols/min/g liver protein and LFD mouse livers consumed 7.04 ± 1.80 µmols/min/g liver protein (Fig. 2H). These results confirmed the accuracy of the HDO analysis to monitor fatty acid β-oxidation from [D_15_]octanoate.

The rate of ketogenesis was also measured from the effluent perfusate for comparison to prior methods of inferring β-oxidation rates as well as for metabolic modeling efforts. The HFD group had a ∼2 fold higher rate of total ketone production than the LFD group (0.85 ± 0.14 vs. 0.41 ± 0.13 μmoles/min) (Fig. 2G). However, these rates were not significantly different when normalized for total liver protein (Fig. 2H), indicating that hepatomegaly is the primary source of overall increased ketogenesis. Additionally, gluconeogenesis rates were measured to assess the contributions of unlabeled pyruvate and lactate during perfusion. Total gluconeogenesis rates were insignificantly elevated in the HFD group (p-value = 0.12) compared to the LFD group (Fig. 2G). No trend was observed in relative gluconeogenesis rates (Fig. 2H).

### Metabolic modeling of NAFLD mouse perfused livers reveals increased β-oxidative activity but a decline in β-oxidative metabolism per g liver protein

To assess global oxidative metabolism in the liver using modeling (Fig. 3A), we calculated oxygen consumption during perfusion as the difference in O_2_ from the afferent and efferent perfusate. Despite the increased liver size after HFD, total oxygen consumption of livers from mice on LFD (2.2 µmols O_2_/min) and HFD (2.25 µmols O_2_/min) were similar (Fig. 3B, left). When normalized for liver protein, oxygen consumption was significantly less in livers from mice on HFD (4.2 µmols O_2_/min/g liver protein) than LFD (8.9 µmols O_2_/min/g liver protein) (Fig. 3B, right). Thus, we inferred that the larger livers of the HFD mice had less active oxidative phosphorylation per gram liver protein than the livers of the LFD mice.

**Figure 3.**
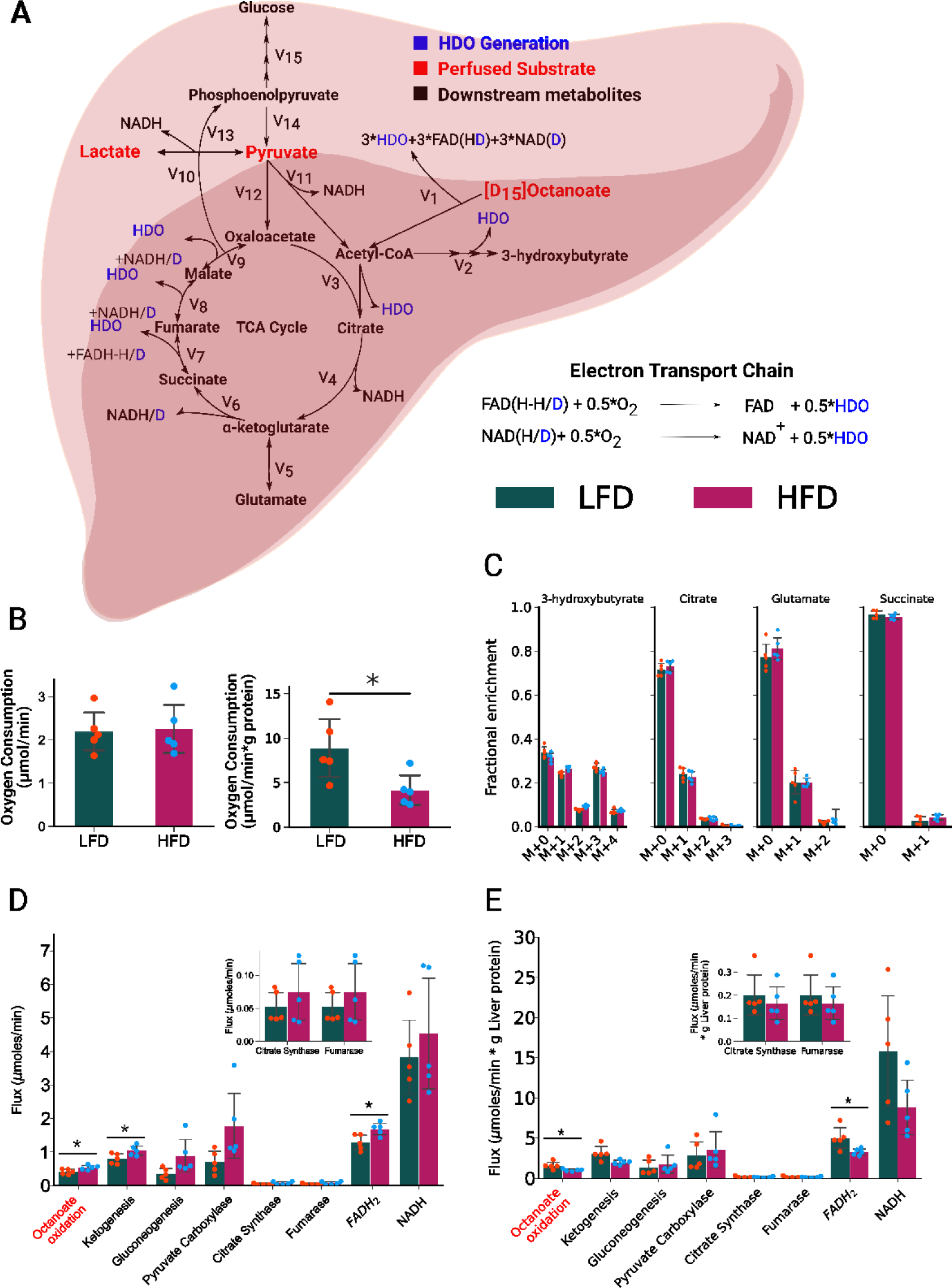
Metabolic modeling of flux in an *ex vivo* perfused liver. (A) Diagram of INCA metabolic model used to define the rates of [D_15_]octanoate consumption, electron transport chain activity and central carbon metabolism. (B) Oxygen consumption was monitored by Hansatech oxygraph+ of afferent and efferent perfusate in [D_15_]octanoate-perfused livers from mice in LFD and HFD groups. Rates of total oxygen consumption and consumption normalized to gram of liver protein are shown. (C) Fractional enrichments of 3-hydroxybutyrate to represent ketones, citrate as an indicator of TCA cycle enrichment and acetyl-CoA partitioned into the TCA cycle, glutamate as an exchangeable portion of the TCA cycle with α-ketoglutarate, and succinate to represent the second half of the TCA cycle exchanging with fumarate. Predicted flux for the mouse perfused liver by INCA metabolic modeling of (D) total flux per liver and (E) relative flux per gram liver protein. Error bars represent mean ± standard deviation. Statistical significance was determined by Student’s t-test for rate comparisons (*, p < 0.05).

To determine specific differences in the oxidative pathways of β-oxidation and the TCA cycle, we built a model with isotopomer network compartmental analysis (INCA) (Fig. 3A) (Table S1) (Fig. S1) [30]. The inputs to the model were as follows: the concentration and isotopologue enrichments of citrate as an indicator of acetyl-CoA incorporated into the TCA cycle; glutamate to capture exchange with α-ketoglutarate; succinate to estimate exchange of 4-carbon TCA cycle intermediates; the rate of ketogenesis, determined by 3-hydroxybutyrate production; oxygen consumption; and the rate of gluconeogenesis. We used GC-MS to measure the pool sizes and isotopomer distribution of metabolites in liver tissue samples and in the effluent of livers simultaneously perfused with [D_15_]octanoate and natural abundance pyruvate and lactate. Of the metabolite pools, only liver-derived alanine showed a significant difference between HFD and LFD mice when normalized for liver protein (Fig. S2). The fractional enrichment of deuterium label in 3-hydroxybutyrate, citrate, glutamate, and succinate across all isotopomers was similar in livers from either HFD or LFD mice (Fig. 3C)(Fig. S3). The estimated acetyl-CoA enrichment of ∼35.5 – 38% was conserved among livers from both groups and was extrapolated from citrate enrichment after adjustment for the loss of one-third of deuterium label resulting from M+1 acetyl-CoA condensation with oxaloacetate. Ketone-extrapolated acetyl-CoA enrichment agreed with this measure after adjustment for multiple labeling by two acetyl-CoA units. Adjustment was achieved by halving 3-hydroxybutyrate enrichment and correcting for the loss of one-third of deuterium label resulting from M+1 acetyl-CoA condensation with acetoacetyl-CoA. The resulting flux estimates demonstrate a clear perturbation of oxidative metabolism (Fig. 3D). Average 95% confidence intervals for parametric optimization of flux (Table S2) demonstrates that elevated oxidative metabolism is consistently congruent with the data. This trend was preserved for the confidence intervals of flux estimates between perfused livers (Table S3).

Both the TCA cycle and β-oxidation produce nicotinamide adenine dinucleotide (NADH) and flavin adenine dinucleotide (FADH_2_) (Fig. 3A, Fig. S1). These two sources of reducing equivalents can be oxidized by the oxidative phosphorylation complex for ATP production. We observed a significantly higher total β-oxidative flux (octanoate oxidation) and FADH_2_ turnover in livers of the HFD group than of the LFD group (Fig. 3D). When normalized for the differences in liver protein, the livers of the HFD mice had less efficient oxidation of octanoate and significantly reduced FADH_2_ oxidation with a trend towards lower NADH oxidation (Fig. 3E, right). The reduced FADH_2_ and NADH oxidation in the HFD mouse livers per gram liver protein provide an explanation for the observed decrease in O_2_ consumption per gram liver protein.

### NAFLD mouse livers exhibit similar abundance of most proteins or transcripts of proteins involved in oxidative metabolism

To determine if there were differences in relative amounts of proteins involved in oxidative metabolism in the livers of mice on the two diets, we measured protein abundance of complex 1 – 5 for the mitochondrial electron transport chain, CPT1, and long chain acyl-CoA dehydrogenase (LCAD) (Fig. S4 A,B). We observed no significant differences in the amounts of these proteins between livers from HFD and LFD mice (Fig. S4 A,B). Additionally, we measured mRNA transcript abundance by qPCR of *LCAD* and *HMG-CS2*, encoding hydroxymethylglutaryl-CoA synthetase 2 (Fig. S4 C,D). *HMG-CS2* mRNA was significantly lower in livers from the HFD animals than those from LFD (Fig. S4C). These data indicated that the source of dysfunction driving differences in β-oxidation was not aberrant protein expression of fatty acid transporters or dehydrogenases.

### DMRI detects hepatic fatty acid oxidation *in vivo*

To assess whether [D_15_]octanoate can serve as a contrast reagent for assessing liver-specific metabolism, we performed deuterium MRI (DMRI) with 12-week-old C57BL/*6J* mice administered [D_15_]octanoate by tail vein injection (Fig. S5). By 24 minutes after [D_15_]octanoate administration, the deuterated signal localized to the liver and was visible by ^2^H FLASH (Fig. 4A-C, yellow circle). Two-point Dixon (2PD) imaging, 52 minutes post injection, indicated that most of the deuterated signal was from HDO (Fig. 4B).

**Figure 4.**
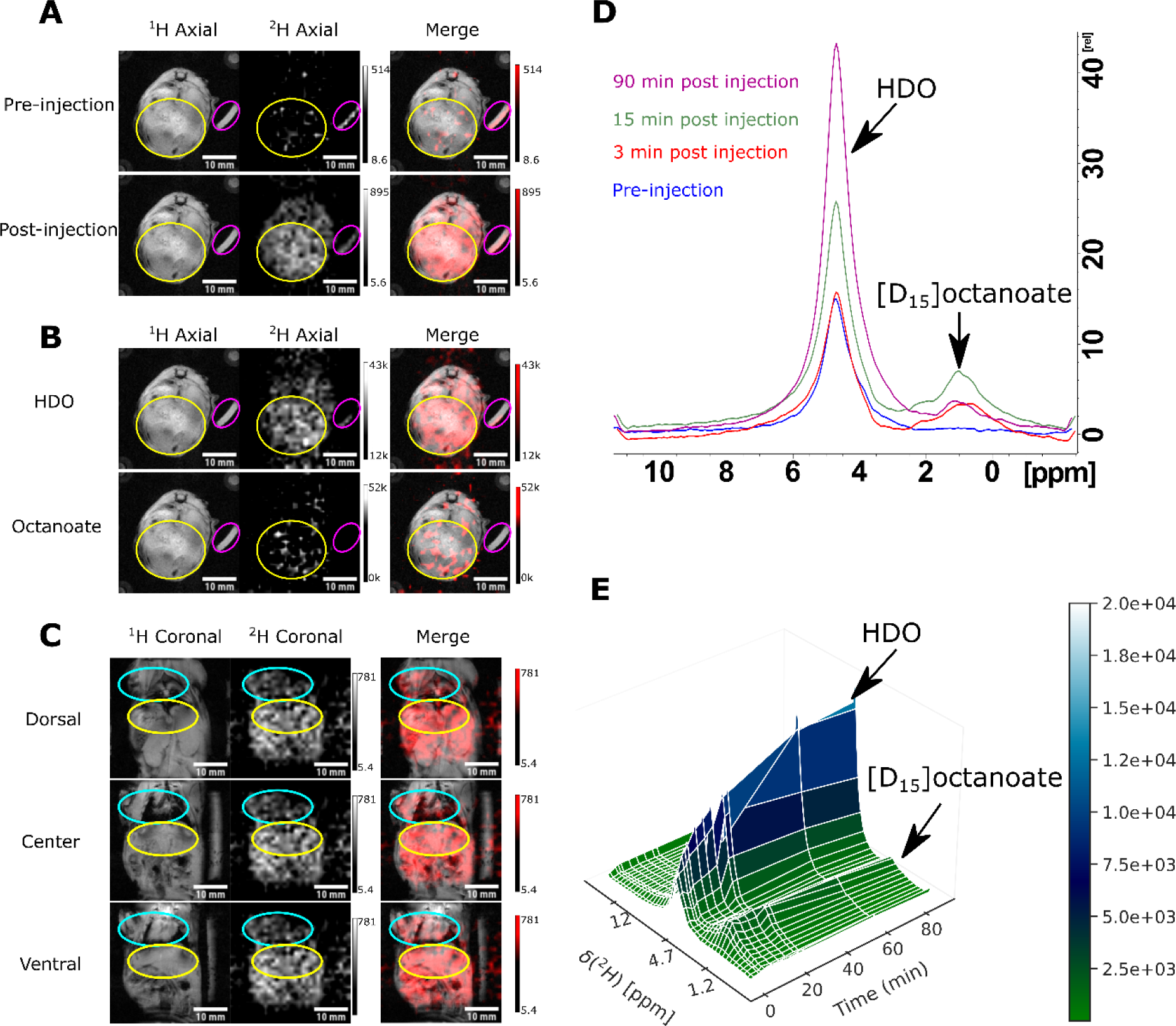
Metabolic characterization of *ad libitum* fed 12-week-old C57BL/6J mouse torso through magnetic resonance imaging and spectroscopy. Images are represented using a ^1^H volume coil (column 1 panels A-C) and a ^2^H saddle coil (column 2 panels A-C) with ^1^H (grayscale) and ^2^H (red) merges of the two (column 3 panels A-C). In panels A-C the liver is circled in yellow and the 0.05% D_2_O standard is circled in magenta. Axial FLASH images through the mouse vertical cross section are taken with respect to the liver in ^1^H and ^2^H modes. The images in panel A represent the signal prior to (pre-injection) and 24 minutes post (post injection) [D_15_]octanoate administration. Overlays of the ^1^H and ^2^H axial images demonstrate localization of the signal to the liver after administration of [D_15_]octanoate tracer. Representative 2PD images obtained 52 minutes after [D_15_]octanoate injection are shown in panel B. In panel C, for the coronal images taken 80 minutes post-injection, the heart and shoulder are circled in teal and the liver is circled in yellow. (D, E) Spectral analysis of fed mouse torso post tail vein injection of [D_15_]octanoate. Data shown represents a single representative fed mouse. In panel D, the spectral overlay of whole-volume, deuterium, single-pulse acquisition pre-injection and 3, 15, and 90 minutes post injection colored with blue, red, green, and magenta respectively. Panel E shows a surface plot of the continuous spectral timeline of deuterium signal: The 4.7 ppm signal corresponds to HDO and the 1.2 ppm signal corresponds to [D_15_]octanoate.

To confirm that the localized signal was primarily from an increase in the HDO signal, we also performed spectroscopic analysis of the torso of fed mice administered [D_15_]octanoate while in the 11 T MRI magnet (Fig. 4D, E). We observed a time-dependent shift in the [D_15_]octanoate and HDO signals (Fig. 4D, Fig. S6). The [D_15_]octanoate signal peaked at ∼5 minutes post-injection and remained stable for 25-30 minutes after injection before slowly declining until 60 minutes post injection. The HDO signal rapidly increased from 0 to 30 minutes followed by a slower rate of increase before peaking at ∼60 minutes after injection (Fig. 4E, Fig. S6).

### DMRI detects dysregulated β-oxidation in NAFLD mice

Having established that we successfully monitored fatty acid β-oxidation by DMRI using [D_15_]octanoate as the contrast label, we tested if we could detect changes in the rate of β-oxidation between *ad libitum* fed and fasted mice in a progressive NAFLD mouse model. We performed DMRI of C57BL/*6J* mice on HFD for 0, 8, 17, and 24 weeks and compared them to mice on a normal chow diet (Fig. 5A, B). We monitored body weight and calorie consumption for both groups of mice, which showed that the HFD mice consumed more calories and had greater overall mass compared to the mice with *ad libitum* access to a normal chow diet (Fig. S7A). The rate of weight gain by the mice with *ad libitum* access to normal chow was stable throughout the experiment, whereas the mice with *ad libitum* access to HFD showed a faster rate of increase for the first 8 weeks and then a rate similar to that of the mice on the normal chow diet. Based on histological assessment of steatosis, the HFD animals in both the fed and fasted groups had NAFLD from 8 weeks of diet onward (Fig. 5C; Fig. S7B, Table S4). Overnight fasted animals on HFD and chow diet had similar liver triglyceride content, potentially a side effect of increased lipolysis and sourcing of extrahepatic fat to the liver for processing and ketogenesis (Fig. S8, left). However, we estimated greater than 5% of hepatic weight was comprised of triglycerides in HFD *ad libitum* fed animals from 8.5 weeks onward (Fig. S8 right, Table S5).

**Figure 5.**
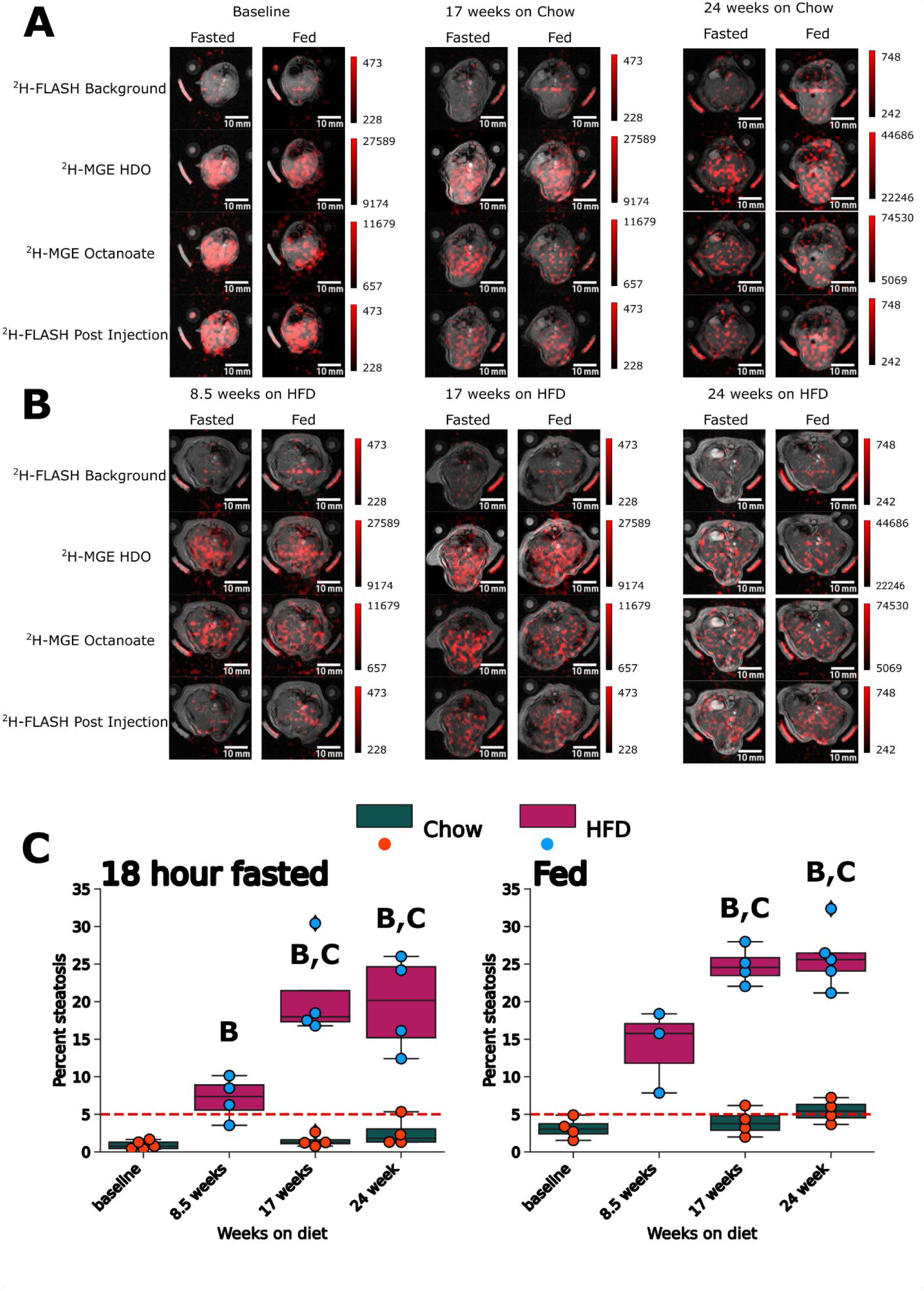
Representative metabolic imaging of C57BL/6J mouse liver for the dietary timeline from 8 weeks of age to 32 weeks of age. All images represent an overlay of the respective deuterium image with the proton axial for the hepatic region. Overlays are ordered from top to bottom per panel as follows: ^2^H-FLASH before injection of [D_15_]octanoate, 27-minute 2PD acquisition starting at 17 minutes post injection (HDO and [D_15_]octanoate components) and 27-minute ^2^H-FLASH acquisition starting at 62 minutes post injection. For each given time point the 18 hour-fasted mice are in the left column and the *ad libitum*-fed mice are in the right column. Images were arranged with respect to timeline in the order of A) Baseline, 17 weeks, and 24 weeks on chow diet. B) 8.5 weeks, 17 weeks, and 24 weeks on HFD. C) Percent steatosis score in 18 hour-fasted (left) and *ad libitum*-fed (right) C57BL/6J mice on either HFD (pink bar, blue dots) or chow diet (green bar, orange dots). The 5% steatosis cutoff for NAFLD is represented as a red dashed bar. Error bars represent the mean ± standard deviation. B refers to significance relative to baseline. C refers to significance when comparing the HFD mice to their chow diet counterparts for the same age. F refers to significance when comparing fed and fasted animals within the same diet and age.

Consistent with the increase in liver mass that we observed for mice on 16 weeks of HFD (Fig. 1B), the imaged livers appeared larger in the mice on the HFD compared with those on normal chow (Fig. 5A, B). We detected an HDO signal following [D_15_]octanoate injection in both fed and fasted control mice on a normal chow diet as well as mice on the HFD (Fig. 5A, B, Fig. S9). We noted that the HDO signal became more punctate and less uniformly distributed in both the HFD and normal chow-fed mice (Fig. 5B), suggesting that metabolism changed as the mice aged during the course of the experiment. This highly intense, punctate HDO signal was notable in both the fed and fasted state at 24 weeks of HFD.

To account for changes in liver size during disease progression, we quantified liver mass for mice in the fasted or fed state at baseline and 8.5, 17, and 24 weeks on each diet (Fig. 6A, Table S6). Comparing liver mass in the fed state confirmed that chow diet mice had significantly increased liver mass at 17 and 24 weeks compared to the mass at baseline (Fig. 6A, Table S7). Liver mass of fed mice was significantly increased after 8 weeks on HFD, compared to baseline. By 17 weeks on HFD, liver mass increased profoundly from 1.22 ± 0.10 g at baseline to 3.45 ± 0.41 g and was significantly greater at both 17 weeks and 24 weeks than that of the mice on normal chow at the same times (Fig. 6A, Table S7). Liver mass of fasted mice was unchanged throughout the experimental period for mice on the chow diet and at 8.5 weeks of HFD (Fig. 6B). At 17 and 24 weeks of HFD the liver mass of fasted mice (2.76 +/-0.58, 2.3+/-0.77 g respectively) was significantly higher than baseline (1.01 +/-0.05 g) or chow diet controls at 17 and 24 weeks (1.36 +/-0.23 g, 1.44 +/-0.28 g) (Fig. 6B).

**Figure 6.**
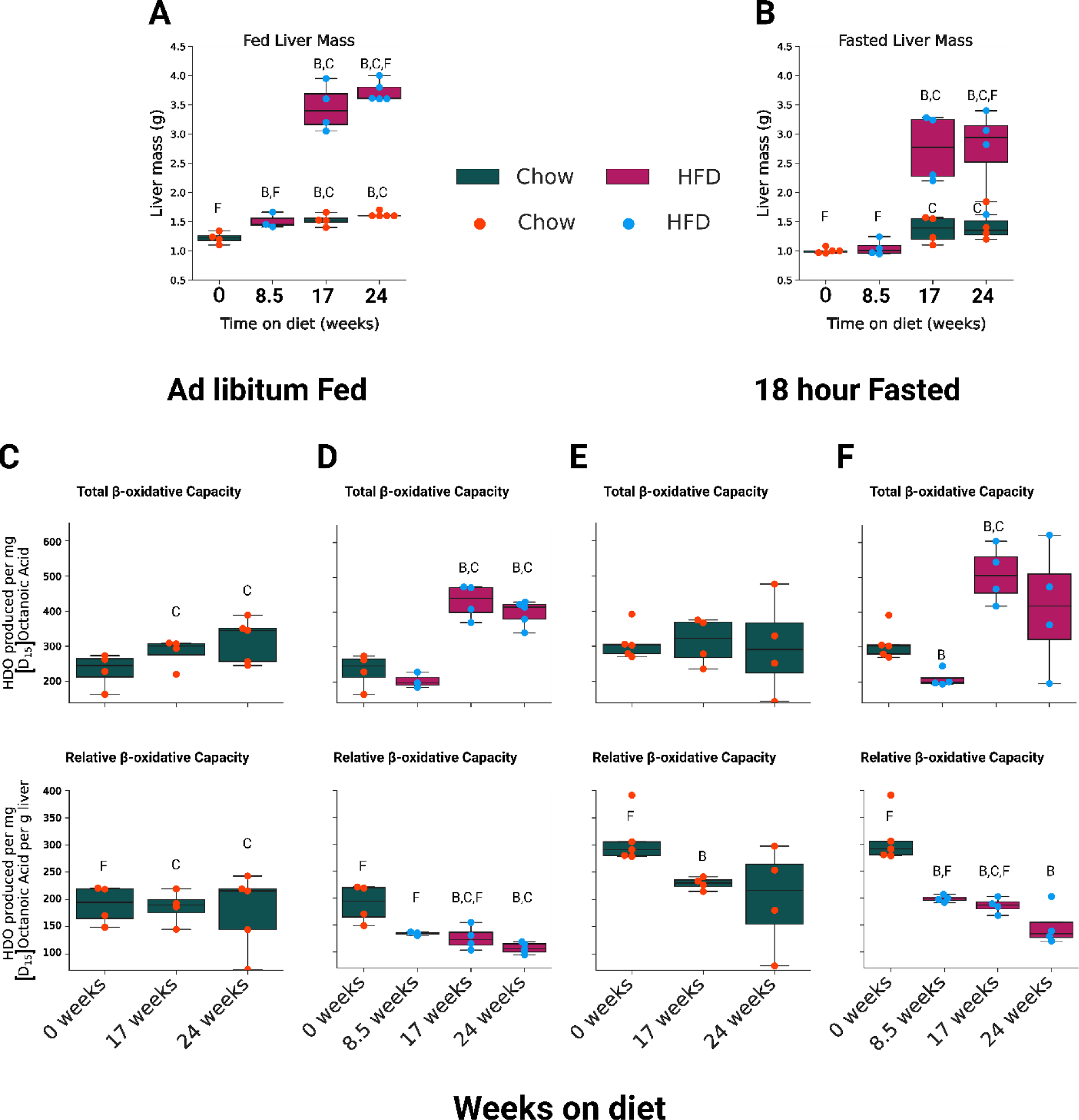
Quantitative measurements of liver mass as well as total hepatic β-oxidation and hepatic β-oxidation relative to liver mass. HFD fed mice are in pink, while baseline and chow fed mice are in dark green. A) Liver mass in grams from ad libitum fed mice or B) 18 hour fasted mice that underwent deuterium MRI at the start of the diet (8 weeks of age) until the end of imaging (32 weeks of age). Weeks on diet was determined by access to the MR imaging system. C-F) MRI quantification of 2PD derived HDO produced per mg [D_15_]octanoate injected (top) and HDO produced per mg [D_15_]octanoate and g liver mass (bottom). See Table S7 for statistical evaluation. The quantification of liver images is ordered as follows: ad libitum fed mouse groups on C) chow diet or D) HFD followed by 18 hour fasted mouse groups on E) chow diet or F) HFD. Error bars represent the mean ± standard deviation. B refers to significance relative to baseline. C refers to significance when comparing the HFD mice to their chow diet counterparts for the same age. F refers to significance when comparing fed and fasted animals within the same diet and age.

We determined both total hepatic β-oxidation and β-oxidation relative to liver mass (see Methods for details) (Fig. 6C-F, Table 1). From baseline through 24 weeks, neither total nor relative hepatic β-oxidation differed in mice with *ad libitum* access to normal chow (Fig. 6C & E, Table S7). In contrast, mice with *ad libitum* access to HFD exhibited a significant increase in total β-oxidative capacity that coincided with the significant increase in liver mass at 17 and 24 weeks (Fig. 6D, top; Fig. 6A). However, when normalized for the increase in liver mass, relative β-oxidative capacity for the HFD mice decreased at all time points compared to baseline (Fig. 6D, bottom).

We found that fasted mice had reduced relative hepatic β-oxidation in mice on normal chow diet that was significant for the 17-week point compared to baseline (Fig. 6E, bottom). At 8.5 weeks on the HFD and following overnight fasting, liver mass was the same as liver mass at baseline of fasted mice (Fig. 6A). However, livers of the fasted mice showed a significant reduction in both total and relative hepatic β-oxidative capacity at this timepoint (Fig.6E). Like we observed for the fed state, livers from the fasting mice on the HFD at 17 and 24 weeks showed an increase in total hepatic β-oxidative capacity but a decrease when normalized for the increase in liver size (Fig. 6F).

## Discussion

### DMR of the perfused liver

Imaging hepatic energy metabolism is not an easily achieved goal, because positron emission tomography (PET) methods depend upon access to a cyclotron for production of ^11^C or ^18^F, and MR-based methods to date have used single-voxel spectroscopy with administration of a large, limiting quantity of ^13^C-labeled substrate. We postulated that imaging with DMR could provide metabolic contrast in a more accessible paradigm than either the PET or previous MR approaches. We first assessed the capacity of monitoring HDO production with DMR to identify changes in β-oxidation for perfused livers. Using a perfused liver model, we removed contributions from extraneous tissues and our results were solely indicative of hepatic metabolism. The [D_15_]octanoate substrate used for the *ex vivo* perfused liver experiments generated a quantifiable measure of β-oxidation through the production of HDO (Fig. 2C,E,F) [23,-31-32]. NMR approaches of this nature are quantitative and revealed that β-oxidation was decreased per g liver protein in the HFD mice, which was confirmed by independent direct measurements of [D_15_]octanoate consumption by GC-MS analysis of perfusate (Fig. 2 G,H). Clearance of [D_15_]octanoate from the perfusate is a similar measure to ^13^C tracer turnover measurements during *in-vivo* assessment of lipid metabolism in NAFLD patients [33–39].

Another common method for inferring β-oxidation with hepatic specificity is the measurement of labeled ketones after tracer administration [40]. β-oxidation produces acetyl-CoA that is required to make ketones (Fig. S1, Table S1), and the liver is considered the primary source of ketogenesis [41]. Especially when coupled with fasting conditions, increased β-oxidation also leads to more ketogenesis, hence analyzing labeling in ketones is frequently employed [40]. Because there was no difference in ketone isotopic labeling pattern for the HFD and LFD groups, we compared total ketone output (Fig. 3C). However, if we had relied on the commonly used metric of ketogenic output alone to determine oxidative rates from the liver, our results would indicate either higher total oxidative potential or no relative difference between dietary states (Fig. 2 G,H) [18–21]. In the perfusion model, where the liver is the only organ, the [D_15_]octanoate consumption from perfusate is a more accurate proxy for β-oxidation than the downstream pathway of ketogenesis and is in complete alignment with our HDO measure by DMR. DMR of HDO produced from deuterated fatty acids therefore bypasses the inherent limitations of less direct β-oxidation measurements.

A limited set of equations modeling only octanoate consumption and HDO production in the perfused liver would produce estimates of β-oxidation with low accuracy due to the uncertainties associated with the multiple pathways converging at acetyl-CoA. A more complete INCA model of central metabolism considerably reduced the error in rate assessments [30]. Our enhanced INCA model of perfused liver metabolism used HDO production, O_2_ consumption, ketone production, and pool sizes and enrichments of multiple metabolites. In addition to validating the claims from our DMR method, we also determined that a decrease in FADH_2_ oxidation in the HFD mouse liver occurred, which has been previously observed in models of β-oxidative decline with isolated mitochondria [42]. NADH turnover trended lower but the lack of significant differences for NADH turnover are likely related to the difference in model variability associated with FADH_2_ metabolism (two reactions) versus NADH in 8 reactions (Table S1) (Fig. 3D,E). The rate estimates from the INCA model also showed significant increases in total β-oxidation and FADH_2_ oxidation for HFD livers, which was in line with ^11^C palmitate PET analysis of oxidative byproducts and isolated mitochondria experiments [21]. These observations support the robustness of using DMR to assess changes in fatty acid oxidative metabolism post injection of [D_15_]octanoate by monitoring HDO. In conjunction with the decreased HDO production rate per gram of liver protein, these results indicated a possible defect in β-oxidative efficiency caused by deleterious effects of HFD, which is consistent with a 16-week HFD study with C57BL/6J mice that showed reduced transcripts for long and medium chain acyl-CoA dehydrogenase (LCAD & MCAD) [20]. Here, *LCAD* expression was insignificantly lower for HFD livers and Western blot analysis showed that protein abundance was unchanged, compared with livers from LFD mice (Fig. S5). This difference in LCAD results with those in the previous study [20] may relate to the greater weight gain exhibited by the HFD animals in our study, which were heavier by 15%, suggesting that the stage of NAFLD was more advanced and the mice had accrued greater damage to the hepatocytes than previously [20].

Unlike the previously used tracer methods, octanoate induces β-oxidation and ketogenesis [18-21, 39-40, 43-48]. As a direct consequence of the higher β-oxidation in our system, octanoate β-oxidation represented a majority of the oxidative metabolism, suppressing the TCA cycle’s contribution (Fig. 3D,E). Emphasis on β-oxidation by this approach is a potential source of differences to other measures of total oxidative metabolism in rodent NAFLD models but may aid in identifying perturbations in the β-oxidative pathway specifically.

### Imaging hepatic oxidative flux

Using HDO as a readout of hepatic β-oxidation has distinct advantages. HDO can be directly imaged, unlike thermally polarized ^13^C methods, which are restricted to a single voxel covering the whole liver in humans. Additionally, based on studies in isolated mitochondria, HDO is retained in hepatic mitochondria post [D_15_]octanoate incubation for at least an hour (Fig. S10). Retention of HDO in liver mitochondria on this time scale suggests that our estimates of hepatic β-oxidation *in vivo* are organ specific. When performing deuterium metabolic imaging, we observed the highest signal in the liver for axial and coronal images relative to the kidneys, heart, and shoulder muscle (Fig. 4A-C). [D_15_]octanoate was clearly used as an oxidative substrate, as evidenced by the rapid increase in HDO pool size detected by deuterium spectroscopy of the torso and 2PD imaging of the liver (Fig. 4A, D,E). Various characteristics of the liver, such as its fenestrated blood vessels, proximity to the heart after tail vein injection, mitochondrial abundance, and physical size, support its major role in β-oxidative metabolism. These characteristics lead us to the conclusion that our *in vivo* estimates of hepatic β-oxidation are organ specific without deleterious contribution from other tissues. Additionally, [D_15_]octanoate is oxidized in the liver in a CPT1 and CPT2 independent manner unlike in mitochondria from skeletal muscle or cardiac muscle [27].

Deuterium metabolic images at 8, 17, and 24 weeks of HFD showed a high degree of signal localization to the liver after injection of [D_15_]octanoate (Fig. 5). Before injection of [D_15_]octanoate, the body noise was very low and the external reference phantoms are the primary component visible in the ^2^H-FLASH images (Fig. 5). At the concentrations used here, no animals suffered respiratory dysfunction post injection. Less signal from the octanoate component in 2PD was observed in the fed than fasted animals (Fig. 5). The signal differences suggested lower amounts of [D_15_]octanoate were taken up by the liver in the fed state, which may contribute to the higher total and relative β-oxidation observed in the fasted animals at multiple timepoints (Fig. 5). As animal size (and age) increased, the distribution of deuterium signal over the liver post injection became less uniform (Fig. 5). The decrease in uniform signal indicated changes in HDO production through less β-oxidative activity, differences in zonation, reduced transport of [D_15_]octanoate, or a combination thereof, throughout the liver. This effect was also observed in the decrease of relative β-oxidation as the HFD progressed (Fig. 6C-F bottom row).

Throughout the course of the HFD, relative β-oxidation decreased most dramatically within the first 8 weeks of HFD and continued to decline in the latter weeks of diet (Fig. 6D/F bottom row). The decrease in relative β-oxidation occurred before major increases in liver mass and total β-oxidation (Fig. 6A,B, D/F top row). The increase in percent steatosis by histological assessment to NAFLD status for HFD animals alone occurred by 8 weeks of HFD (Fig. S7, S8). The timeline implied that excess lipids and higher caloric intake in the HFD led to a decrement in liver β-oxidative capacity before major changes in hepatic structure towards hepatomegaly and greater total hepatic β-oxidation occurred. Unlike HFD animals, chow diet animals did not have a significant increase in total β-oxidation (Fig. 6C/E top row). A small decrement in relative β-oxidation for overnight fasted chow diet animals was observed at 17 weeks of diet only (Fig. 6E bottom row). These results confirmed that changes in physiology caused by the HFD are detected with this method. Therefore, DMRI after injection of [D_15_]octanoate is a powerful tool for assessing hepatic β-oxidation directly, with a limited contribution from TCA cycle metabolism. Although octanoate administration downregulates TCA cycle turnover, we propose that the metric of β-oxidation produced by these methods can generate insights into dysregulated hepatic metabolism, such as those induced by HFD. DMR post [D_15_]octanoate administration is quantifiable and metabolically sensitive and can safely monitor NAFLD progression *in vivo* in a mouse model of the human disease. Because DMRI is already practiced in humans with deuterated glucose, this approach should be readily translated to human studies.

## Conclusion

We demonstrated that HDO production from deuterated fatty acids is a viable methodology for estimating functional differences in the rate of hepatic β-oxidation in a diet-induced model of fatty liver disease. β-oxidation inactivity has been cited as a contributor to excess fatty acids that drive ER stress and lipoapoptosis [15–17]. Conversely, increased β-oxidation, in a high fatty acid environment during NAFLD, acts as a direct contributor to the overactive production of inflammatory molecules such as ROS [18–22]. We directly measured hepatic β-oxidation *ex vivo and in vivo* with DMR of hepatic deuterated fatty acid metabolism NAFLD (Fig. 2D,E,F,H, Fig. 3D&E, Fig. 6C-F). Our results suggested that both views of fatty acid oxidation are valid. Whole liver fatty acid metabolism is increased in this model, but the oxidative process is less efficient in the diseased liver. The sensitivity of this methodology, which is inherently non-invasive, provides an attractive route for new paradigms of clinical assessment for NAFLD diagnosis, monitoring, and treatment.

## Methods

### Liver perfusion

Experiments involving mice (strains used were C57BL/6*J* (Stock No. 000664)) were handled in compliance with University of Florida Institutional Animal Care and Use Committee (protocol number #201909320). All male mice were 8 – 10 weeks old at the start of the diet. Mice were maintained on either a LFD (10% Kcal from fat, D12450J, Research Diets, New Brunswick New Jersey) or HFD (60% Kcal from fat, D12492, Research Diets New Brunswick New Jersey) for 16 weeks and were weighed every week. At 16 weeks on the diet, mice were overnight fasted 12-14 hours before being anesthetized using isoflurane followed by an intraperitoneal injection of 200 μL of saline with 200 units of heparin. Approximately ten minutes after heparin injection, a celiotomy was performed under anesthesia. Lidocaine was administered subcutaneously prior to making the first incision that exposed the liver and the portal vein. The portal vein was then cannulated, and perfusion was started. The liver was then excised from the body and connected to the glass perfusion column [49], which was moved into the bore of a 600 MHz NMR magnet.

Livers were perfused with Krebs – Henseleit buffer (25 mM NaHCO_3_, 112 mM NaCl, 4.7 mM KCl, 1.2 mM MgSO_4_, 1.2 mM KH_2_PO_4_, 1.25 mM CaCl_2_, and 0.5 mM sodium-EDTA), 1 mM sodium lactate, and 0.1 mM sodium pyruvate during magnet shimming. The system was switched to recirculating mode after magnetic field shimming (∼12-16 minutes), and the perfusate solution was supplemented with 1.2 mM sodium-[D_15_]octanoate. Perfusate was oxygenated using 95% O_2_/5% CO_2_ mixed gas (Airgas) for the duration of perfusion. Oxygen consumed by the liver during perfusion was measured every ten minutes using an Oxygraph+ setup (Hansatech Instruments, UK). At the end of perfusion, the livers were freeze clamped using liquid nitrogen and stored at −80°C. One mL perfusate samples were collected every 10 minutes after switching to the [D_15_]octanoate perfusate for analysis of HDO production and GC-MS metabolomics. The number of mice used in each group were LFD mice (n = 5) and HFD mice (n = 5).

### NMR spectroscopy

Deuterium spectra were collected in a custom built 20 mm broadband probe (QOneTec, Switzerland) installed in a 14 T (92 MHz ^2^H frequency) magnet equipped with an Avance III NMR console (Bruker Biospin, USA). Deuterium spectral acquisition began one minute prior to switching to perfusate with the [D_15_]octanoate. Magnetic field shimming was executed by manual adjustment while observing the linewidth of the ^23^Na signal associated with the perfusate. Typically, linewidths of around 18 Hz in ^23^Na spectrum were achieved with ^2^H spectral linewidths also averaging ∼18 Hz. ^2^H spectra (spectral width of 25.0192 ppm; 4096 data points) were recorded using a 90° pulse centered on the HDO resonance (nominally, 4.7 ppm) with ^1^H decoupling (WALTZ65; B1 = 4.5 KHz) during the acquisition (acquisition time=0.887 s) and a repetition time of 4.9 s with 16 scans per 1.33 min time point and up to 128 time points. Spectra were processed with 1.5 Hz exponential line broadening and baseline corrected using a polynomial function. Peak areas of HDO and octanoate were obtained by integration of the 5.4 to 3.8 ppm and 2.4 to −0.3 ppm regions, respectively, with a custom python script.

### Gas chromatography – mass spectrometry

#### Perfused liver extraction

Freeze-clamped samples of the perfused liver were stored at −80 °C prior to analysis. Perfused liver samples (110 ± 10 mg) were transferred into 2 mL microcentrifuge tubes with screw caps, and 1.0 mm zirconium oxide homogenization beads were added along with 1 mL of cold degassed acetonitrile:isopropanol:water (3:3:2 v/v/v) solvent mixture. Samples were homogenized with a bead homogenizer (Fastprep-24, M.P. Biomedicals, Irvine, CA) for 3 x 20 sec cycles and were cooled on ice for 5 minutes between each cycle. The samples were then centrifuged at 10,000 x g at 4 °C for 30 min. Supernatant (900 µL) was recovered and lyophilized (Thermo Scientific Waltham, MA). The dried precipitate was reconstituted in 150 μL of acetonitrile:water (1:1 v/v) mixture followed by incubation at −20 ° C for at least 2 hours. Reagents, unless otherwise specified, were purchased from Fisher scientific, New Jersey.

### MTBSTFA derivatization

For analysis of amino acids, ketones, and TCA cycle intermediates, ∼8 mg of liver extract or 20 µL of perfusate were dried down by gentle airstream. The sample was reconstituted in 50 µL of methoxyamine HCL in pyridine (Thermo Scientific, Pennsylvania) for 1.5 hours at 30 °C with stirring. N-tert-Butyldimethylsilyl-N-methyltrifluoroacetamide (MTBSTFA; ProteoSpec MTBSTFA w/ 1% TBDCMS, Ricca Chemical Company, Texas) (50 µL) was added and samples were incubated at 70 °C for 30 minutes while stirring. After derivatization, 80 µL of the sample was transferred to a GC vial for analysis by GC-MS.

### Aldonitrile pentapropionate derivatization

For measurement of sugars eluted during perfusion, 500 µL of perfusate was dried in 0.5 mL V-vials and reconstituted in 50 μL of 20 mg/mL hydroxylamine hydrochloride (CAS# 5470-11-1; Acros organics, New Jersey) in pyridine (CAS#25104, Thermo scientific, Rockford, IL) and stirred at 90 ° C for 1.5 hours. Propionic anhydride (CAS# 240311-50G; Sigma Aldrich, St. Louis, MO) (100 μL) was then added and samples were stirred at 70 °C for 30 minutes. Samples were dried by gentle airstream and then reconstituted in 100 µL of ethyl acetate for GC-MS analysis.

### Metabolite analysis by GC-MS

After the derivatization reaction (aldonitrile pentapropionate or MTBSTFA) was completed, 5 μL of each sample was pooled together to make a “pooled sample” for a given derivatization batch and 80 μL of each individual sample were loaded into a 150 μL glass insert (CAS#13-622-207, Thermo scientific, Florence, KY) inside of a GC vial with a 9 mm PTFE/red rubber septum (CAS#C4000-30, Thermo scientific, Rockwood, TN). Samples were placed in an AI/AS 1300 autosampler prior to auto injection. A 1 μL injection volume was used for each sample into the injection port of the Trace1310 Gas chromatography system (Thermo Scientific, PA). The injection was performed in splitless mode with a splitless single taper liner, splitless time of 60 seconds, and injection port temperature of 250 ° C. Column flow was set at 1 mL/min through a 30 m RTX-5MS integra Guard column (Crossbond 5% diphenyl/95% dimethyl polysiloxane CAT# 12623-127, Restek PA) and 10 m guard column. For MTBSTFA-derivatized samples, the following parameters were used: 60 °C hold time = 1 minute, ramp of 15 °C/min up to 320 °C followed by a 5-minute bakeout/cleaning phase at 320 °C. Aldonitrile pentapropionate– derivatized samples were run on the same GC-MS instrument with the following parameters: start temperature of 80 °C with a hold time = 1 minute, 20 °C per minute ramp up to 280 °C, followed by a 5-minute bakeout at 280 °C. For all samples, the transfer line was maintained at 280 °C and the ion source was maintained at 230 °C. The solvent delay on the mass spectrometer was set to 10 minutes for MTBSTFA-derivatized samples and 8 minutes for aldonitrile pentapropionate–derivatized samples. A m/z filter of 40-600 mass-to-charge ratio was used for spectral acquisition. Individual metabolites were identified and quantified by area using their known quantification ions (Table S8). Concentrations were assessed against an eight-point standard curve that ranged from 50 to 1600 ng of the metabolite of interest.

### Peak integration

Peak areas were integrated using Xcalibur (version 4.1) batch processing with genesis peak fitting, a 5-second time interval, height cutoff at 5.5% of the peak with valley detection enabled, and a signal to noise (S/N) cutoff threshold of 3.

### Isotopic ratio and fractional enrichment

The isotopic ratio of each metabolite for its given quantitation ion was calculated as the ratio of intensity of each mass isotopomer (*m_i_*) to the sum of all mass isotopomers. The isotopic ratio distribution was then corrected for natural abundance as previously described using the Isotopomer Network Compartment Analysis (INCA) software [30,50].

### INCA metabolic modeling

A metabolic model (Table S1) was designed in INCA to track production of HDO through β-oxidation and TCA cycle turnover. To provide a boundary condition dependent on total, quantitative, oxidative flux, the model was supplied with measurements of oxygen consumption, fractional enrichment and pool sizes for selected TCA cycle intermediates and the ketone 3-hydroxybutyrate, rates of gluconeogenesis and ketogenesis based on the final concentration in the effluent, and the fold change increase in the HDO peak over the course of the experiment.

The label in citrate derives from deuterated acetyl-CoA generated during β-oxidation in this system (Fig. 3B, Table S1). Ketones are derived from two individual acetyl-CoA units and thus can be labeled if either one of the acetyl-CoA units is labeled. Consequently, the probability of attaining a labeled ketone is nearly double that of the acetyl-CoA fractional enrichment. Thus, it could be estimated that the acetyl-CoA fractional enrichment is 33.5%. However, M+1 generated acetyl-CoA units (Figure S1) have a 1/3 chance of isotopic loss if they act as a nucleophile in the generation of the ketone 3-hydroxybutyrate. If we assume that M+1 generated acetyl CoA units have an equal chance of being an electrophile or a nucleophile in the generation of ketones, then we would expect a loss in ketone enrichment equivalent to 1/6^th^ of the M+1 fractional enrichment (∼4%). Thus, an additional 2% can be added to the overall acetyl-CoA enrichment to yield a 35.5% enrichment. Citrate enrichment is similarly affected by M+1 deuterated acetyl-CoA label loss during nucleophilic attack by a 1/3 ratio. In this situation the M+1 fractional enrichment (23.4%) of citrate is estimated to be 66% of the actual M+1% fractional enrichment. To get an accurate estimate of M+1 acetyl-CoA fractional enrichment, lost citrate M+1 enrichment must be added back in by taking 50% of the current citrate M+1 fractional enrichment. After adjustment for label loss the total enrichment from acetyl-CoA in citrate is estimated to be 38.8%. After correction, the fractional enrichment of acetyl-CoA is ∼ 35-39% (Fig. 3C, D; Table S1). Label in the TCA cycle decreased from citrate (25%) to fumarate (0-2%) with glutamate and succinate at 18% and 8%, respectively (Fig. 3C) (Fig. S3). Based on these fractional enrichments the deuterium label is liberated before the beginning of the second turn of the TCA cycle. Therefore, all label in the TCA cycle represents first pass kinetics.

### Protein estimation in liver tissue

Protein was extracted from 10 mg of powdered excised mouse liver sample by homogenization with 300 μL of 1X RIPA lysis buffer in a bead homogenizer (Fastprep-24, M.P. Biomedicals, Irvine, CA). Samples were mixed by inversion at 4 °C for 2 hours. After mixing, the samples were centrifuged at 10,000 x g at 4 °C for 30 minutes before collecting 150 μL of the supernatant. Samples were diluted 15-fold before using 10 μL of diluted liver protein extract with 200 μL of Bradford reagent and incubating for 5 minutes before quantifying protein absorbance at 595 nm on a 96-well plate against a standard curve of BSA protein standards that ranged from 1.2 to 0.0625 mg/mL.

### Expression profiles in liver tissue

For quantitative reverse transcriptase-polymerase chain reaction (QPCR) analysis, frozen liver tissue (∼10-15 mg) was homogenized in 600 µL TRIZOL reagent (Invitrogen, Carlsbad, CA) to isolate mRNA using a miniprep kit (Bio-Rad Laboratories Inc., Hercules, CA). A cDNA synthesis kit (Bio-Rad, Hercules, CA) was used to prepare cDNA from 1 µg of mRNA. QPCR was performed utilizing 25 ng of cDNA, 150 nM forward and reverse primers, and 5 μl of SYBR green PCR master mix (Invitrogen, Carlsbad, CA) with *cyclophilin* as the housekeeping gene. Each sample was run in triplicates on a Bio-Rad CFX Real-Time system (C1000 Touch Thermal Cycler).

For Western blot analysis, frozen liver tissue (∼ 10-15 mg) was lysed in 1x RIPA lysis buffer with protease and phosphatase inhibitors. The solution was centrifuged at 15,000 RPM (20,627 G) for 15 min at 4 °C. Total protein was analyzed by Pierce BCA protein assay kit (Thermo Fischer Scientific. Waltham, MA). Protein separation by gel electrophoresis was performed using Bolt 8 % Bis-tris Plus gels (Invitrogen, Carlsbad, CA) before transfer to a nitrocellulose membrane. The membrane was incubated with the primary antibody overnight followed by incubation with the secondary antibody. Primary antibodies against CPT1A, COXI-V, GAPDH (Cell Signaling Technology, Danvers MA), and LCAD (Abcam, Cambridge, UK) were utilized for this study. Western blot quantification was performed with imageJ of the respective blots signal relative to GAPDH signal.

### Mitochondrial isolation to determine HDO retention

Whole liver was excised from a 12-week-old Sprague Dawley rat after sacrifice and cut into 1 gram pieces before crushing in a Dounce homogenizer in 15 mL of isolation buffer I (225 mM mannitol, 75 mM sucrose, 10 mM HEPES potassium, 1 mM EDTA, 0.1 % fatty acid free BSA, pH 7.4) [45] and then centrifuging the homogenate at 1300 x g for 30 minutes at 4°C. The supernatant was then collected in a new tube and centrifuged at 10,000 x g for 30 minutes at 4°C. The supernatant was then discarded. The pellet was resuspended in 2 mL of isolation buffer II (225 mM mannitol, 75 mM sucrose, 10 mM HEPES potassium, 0.1 mM EDTA, pH 7.4), transferred to a clean 2 mL centrifuge tube, and centrifuged at 10,000 x g for 30 minutes at 4°C. The supernatant was then discarded and the pellet was resuspended in 200 μL isolation buffer II. 67 µL of mitochondrial pellet was incubated with 2 mL of incubation buffer (125 mM KCL, 2 mM K_2_HPO_4_, 5 mM MgCl_2_, 10 mM HEPES, 10 μM EGTA, pH 7.2) 4mM sodium octanoate or 4 mM sodium-[D_15_]octanoate and 0.2 mM of ADP for 5 minutes at 37°C before taking the buffer and mitochondria and pelleting them out at 10,000 x g for 30 minutes at 4°C. The supernatant was collected for further analysis of HDO enrichment by NMR and the pellet was subjected to 2 more washes of 1 mL of isolation buffer I and two more centrifugations at 10,000 x g fro 30 minutes at 4°C before resuspending the pellet in 300 uL of acetonitrile with a 2.5 mM [D_4_]pyrazine standard to lyse the pellet and free internal water storage for HDO analysis.

### Deuterium Magnetic Resonance Imaging

Initial metabolic images were acquired as in Mahar *et al.* 2021 [24] with the following modifications: a 26 mm deuterium saddle coil was used instead of 14 mm deuterium surface coil, the coil was placed under the animal’s abdomen, and the average time for image collection was 26 minutes instead of 13 minutes. 12 week old male C57BL/*6J* mice were placed in the magnet and a background image was taken. After injection with 0.675 mg/g [D_15_]octanoate in a 180 μL volume through a catheterized tail vein, the imaging protocol continued (Fig. S5). Two point Dixon (2PD) imaging was performed with respect to HDO and [D_15_]octanoate [51–52].

For the MRI diet study, 8-week-old C57BL/6J mice were maintained on a standard rodent maintenance diet (Teklad global protein 18%, with 18% of calories from fat) or a HFD (60% Kcal from fat, Research diets) for up to 24 weeks. Food consumption and animal weight were measured weekly for both mice cohorts. Imaging sessions took place at 0, 8, 17, and 24 weeks of diet. For each imaging session, mice were anesthetized with isoflurane, then weighed and the tail vein was catheterized. Subsequently, the animal was placed in a 26 mm linear deuterium saddle coil while under isoflurane anesthesia and the coil was tuned to −42 to −55 db using a network analyzer. Then the animal was placed within the 11 T, horizontal bore magnet. After centering the mouse’s liver within the coil and shimming down to an HDO peak linewidth of 80 Hz or less, background single pulse, selective pulse fast low angle shot gradient (SPFLASH) of HDO and [D_15_]octanoate internal standards, liver SPFLASH spectra and ^2^H FLASH images were acquired (Fig. S9). Slice thickness was adjusted to span the liver region within an axial slice without overlap with the heart or kidneys (between 8-14 mm). Afterwards the mice were injected with 0.33 mg/g [D_15_]octanoate, with a maximal injection of <14 mg. Ad libitum fed animals were injected at 10:00 a.m. ± 30 minutes and ∼18 hour-fasted mice were injected at 2:00 p.m. ± 30 minutes. Injections were performed at a rate of 40-50 μL per minute with a 0.25 M sodium-[D_15_]octanoate solution pH 7.2. During and after injection, several single pulse, unlocalized spectra and SPFLASHs of the phantom and tissue were obtained to view import and utilization kinetics. The 2PD imaging was acquired for HDO versus [D_15_]octanoate signal from 17 to 44 minutes. The 2PD image was followed by a whole torso slice selective excitation of the HDO phantom first then the [D_15_]octanoate phantom second, and SPFLASH of the liver region last (occurring between 45- and 52 minutes post-injection). This was followed by a full spectral image at 62 to 90 minutes and then a repeat of the spectroscopy that followed the 2PD imaging. Finally, a follow up ^1^H axial image was obtained for co-registration. Deuterium images were overlaid with ^1^H images and the ratio of deuterium signal over the liver region relative to the HDO phantom was used to determine the fold change in average signal before and after [D_15_]octanoate injection. The fold change was then corrected for total signal change by multiplying the fold change by the area of the liver and the depth of the image (total volume). The total fold change was converted into the total signal fold change per mg dose of [D_15_]octanoate injected to accurately assess total β-oxidation per mg dose and total β-oxidative capacity of the liver. An estimate of the β-oxidative efficiency of the liver was calculated by dividing the total β-oxidative capacity by the liver mass.

### Histological assessment of steatosis in diet mice

The medial lobe of the liver located to the right of the gall bladder was collected for histological assessment of liver disease and steatosis. The excised tissue was incubated in a 2% formalin solution overnight at room temperature before transfer to fresh 70% histological grade ethanol the following day and another overnight incubation at room temperature. On day two after collection the medial lobe was transferred to a fresh solution of 70% histological grade ethanol and incubated overnight at room temperature. On day three post collection the tissue, in 70% histological grade ethanol, was placed within a 4°C fridge until it was submitted to the University of Florida Molecular Pathology Core to be paraffin embedded, cut onto a slide and stained via a hematoxylin and eosin (H&E) stain protocol. The stained tissue was imaged within the University of Florida Molecular Pathology Core and images were processed using ImageJ to calculate the percentage steatosis by lipid fraction area. Five subregions of each liver slice were utilized to determine the percent steatosis score. Lipid fraction area was determined by calculating the white space in the liver fraction with a grayscale cutoff of 190 to 255 and manual removal of white space due to arterial and venous fractions of the tissue.

### Liver triglyceride quantification

100 mg of freeze clamped powdered liver tissue was homogenized in 1 mL of IGEPAL CO-630 – average M_n_ −617 (Sigma Aldrich, St. Louis, MO). Samples were heated at 80 °C in a metal bead bath for 5 minutes then cooled to room temperature and this process was repeated twice until the solution was cloudy. The solution was centrifuged for 2 minutes at 10,000 x g at room temperature to remove insoluble material. 100 uL of supernatant was collected and diluted 100-fold with water before assay. 1.5 μL of the diluted sample was added into a wells in a 96 well plate. Samples were raised to a final volume of 50 μL with Triglyceride Assay Buffer (CAS# MAK266, Sigma Aldrich, St. Louis, MO). 2 μL of Lipase was added to each sample and was incubated for 20 minutes at room temperature to convert triglyceride to glycerol and fatty acids with the plate shaken manually every ten minutes. Then 50 μL of master reaction mix was added to each well and the plate was incubated in the dark with occasional stirring every ten minutes for 30 minutes. Subsequently, absorbance at 570 nm was measured by a plate reader . The absorbance was blank corrected against assay buffer and quantified relative to a standard curve of 0.1, 0.4, 1.6, 6.4 and 10 nmol’s per well.

### Statistical analysis

For comparison of dietary timepoints and binary groups students t.tests (two-tailed) were used to establish significance with an α=0.05. As the number of variables was limited, no correction for multiple variables was used. For HDO production and kinetic measurements significance was determined based on analysis of covariance (ANCOVA) via in lab scripts using Rstudio and the in-built aov function.

### Software

All NMR data processing was carried out using Bruker Topspin (3.6.2+) or Mestrenova (*v*14.0.1-23284 (Mestrelab Research S.L., Santiago, Spain). Image processing for DMRI was carried out using in lab designed MatLab (Mathworks, Natick, MA) scripts and ImageJ. Figures were prepared using in house python (v3.8) scripts: Libraries utilized were numpy (v1.19.2), pandas (v1.1.3), matplotlib (v3.3.2), and nmrglue (v0.8), and Inkscape (v0.92.2). Metabolic modeling was performed using INCA v1.9 [30].

