## Supplemental data for "Detecting altered hepatic lipid oxidation by MRI in an animal model of NAFLD"

### Supplementary Data

#### Localization of HDO to the mitochondrial compartment

The results from the perfused liver indicated that HDO was a biomarker of β-oxidation. However, generalization of the method to *in vivo* requires that HDO is generated and retained in the liver, because octanoate is oxidized in many tissues in the whole animal. On the basis of the accumulation of deuterated signal in the livers of mice injected into the tail vein with [$D_{15}$]octanoate (Fig. 4), we hypothesized that HDO does not freely diffuse through the mitochondrial membrane. To provide sufficient material for analysis, we isolated mitochondria from livers of Sprague Dawley rats and assessed β-oxidation of either deuterated or unlabeled octanoate. Hepatic isolated mitochondria were incubated with 4 mM [$D_{15}$]octanoate and subjected to two, 1-mL washes with natural abundance $H_2O$ buffer over the course of 1.5 hours at room temperature to wash out freely diffusing deuterated water. We determined the amounts of HDO released into the buffer and associated with the mitochondrial pellet (Fig. S10). Compared to the HDO concentration in the buffer from mitochondria incubated with unlabeled octanoate, the buffer from the mitochondria incubated with [$D_{15}$]octanoate exhibited an insignificant increase. In contrast, we detected a ~50% increase in HDO in the mitochondrial pellet following incubation in [$D_{15}$]octanoate, thus, confirming that HDO produced by metabolism of octanoate accumulated in mitochondria of rodent livers.

Because mitochondria are the site of β-oxidation and the mitochondrial membranes tightly control $H^+$ flux to enable chemiosmotic ATP production, we hypothesized that mitochondria were the source of the HDO signal that did not rapidly diffuse. Assuming free diffusion of the water, the dilution factor after 2 subsequent washes of 1 mL exceeds 1000-fold. A 1000-fold dilution should return mitochondrial HDO enrichment to natural abundance, as the greatest enrichment observed *in vivo* was only 3-fold natural abundance of HDO (Fig. S5). The incubation buffer measured a nonsignificant ~6% increase in HDO signal from [$D_{15}$]octanoate treated mitochondria compared to unlabeled sodium octanoate (Fig. S10 A), which indicated negligible export of HDO. However, after two washes a ~50% increase in HDO signal for the [$D_{15}$]octanoate-exposed mitochondrial pellet compared to the unlabeled octanoate-exposed mitochondrial pellet was maintained (Fig. S10). If the mitochondria could freely exchange HDO, the added [$D_{15}$]octanoate would have to produce a 20% enrichment in HDO in the mitochondria to produce the observed enrichment after a ~1000 fold dilution. We note that malate, a hydride receptor in the TCA cycle that exists in rapid equilibrium with oxaloacetate, was only ~0-4% enriched in $^2H$ (Fig. S3). If the mitochondrial compartment had a 20% enrichment, malate should reflect this fact. Collectively, these observations showed that diffusion of HDO across the mitochondrial membrane is limited.

### Supplementary Figures

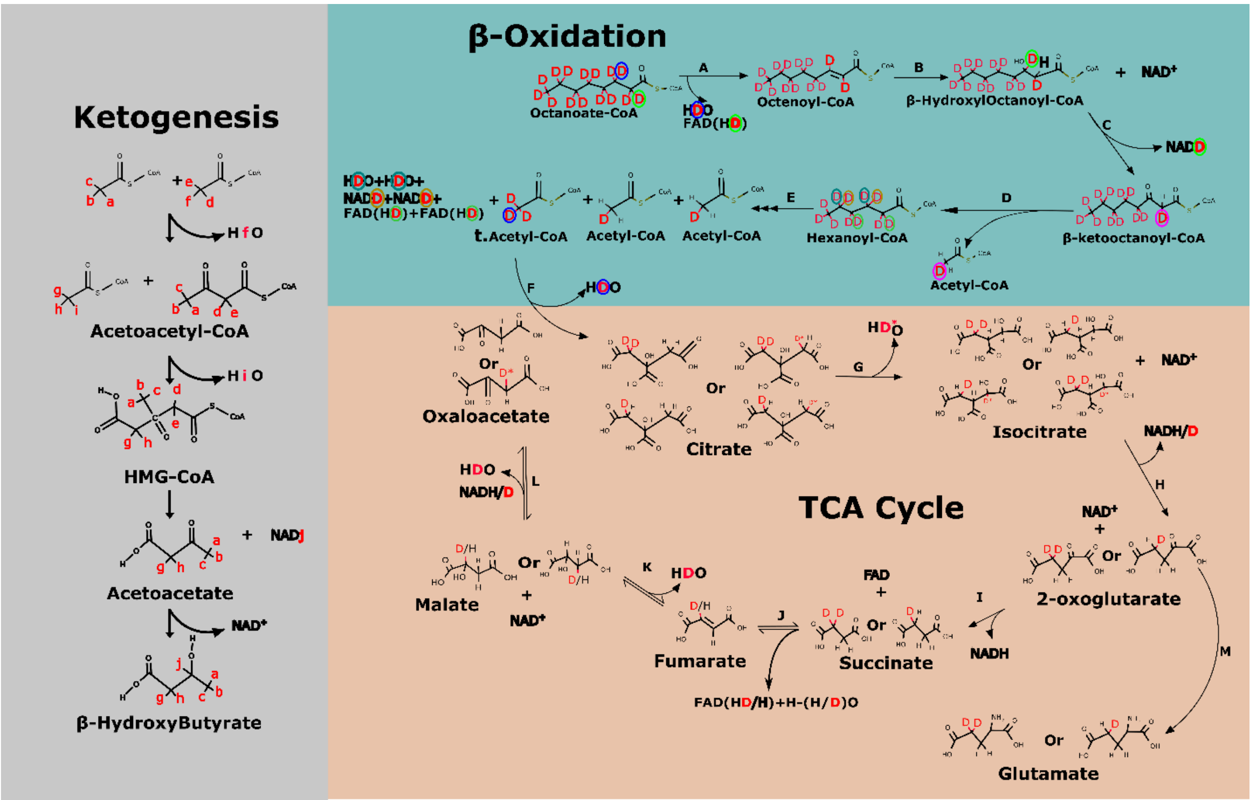


**Figure S1. Model with atom tracking of deuterium elution during fatty acid oxidation.** The pathway for deuterium elution from [$D_{15}$]octanoate starts with β-oxidation. Once acetyl-CoA is produced it can participate in ketogenesis with labeling possible on all red atoms or in the TCA cycle with possible deuterium labeling presented as red D's for deuterium.

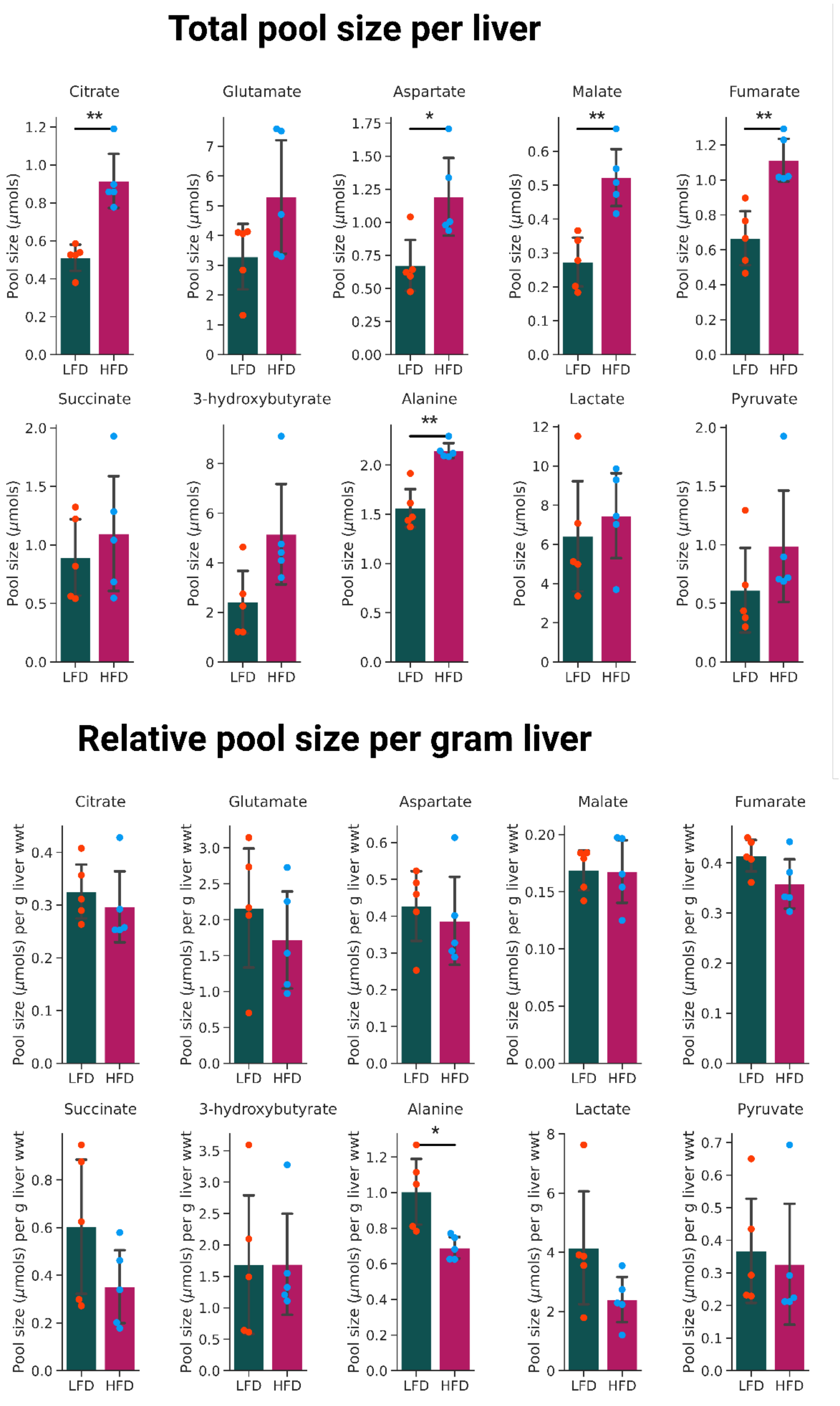


**Figure S2. Total and relative pool sizes of amino acids and TCA cycle intermediates in the perfused liver.** Significance was determined using Student's t-test with *, **, *** corresponding to p-values less than 0.05, 0.01, and 0.001 respectively.

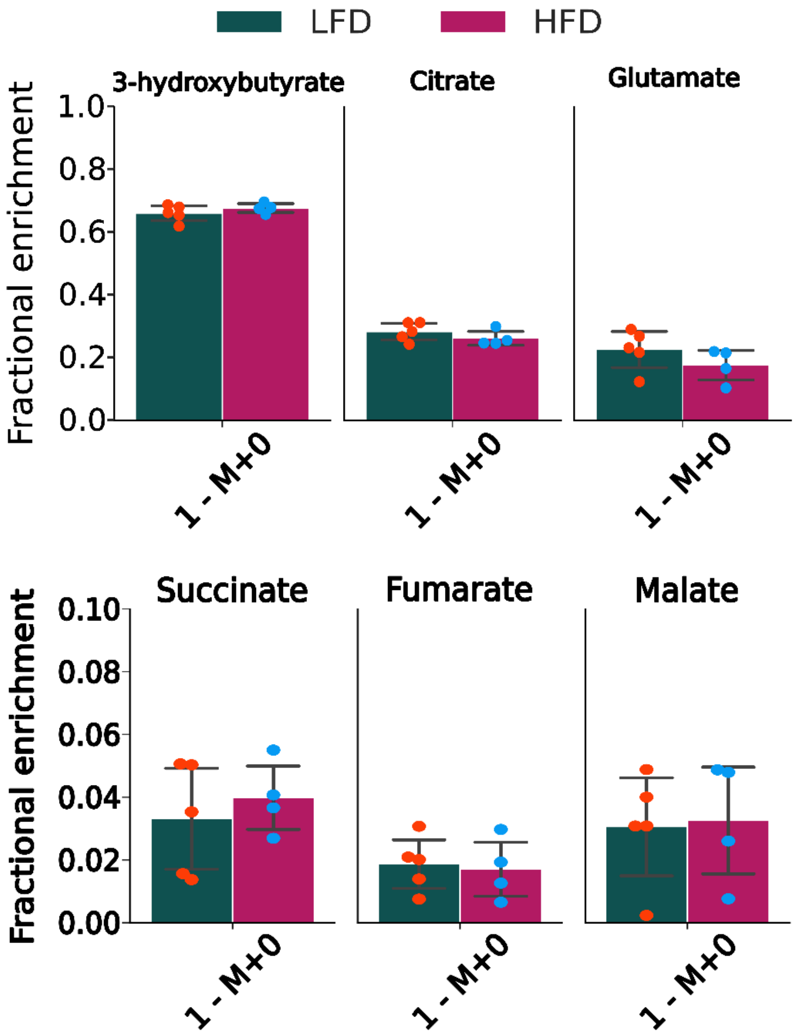


**Figure S3. Fractional enrichment represented by the 1 – M+0 fraction of 3-hydroxybutyrate, citrate, glutamate, succinate, fumarate, and malate.**

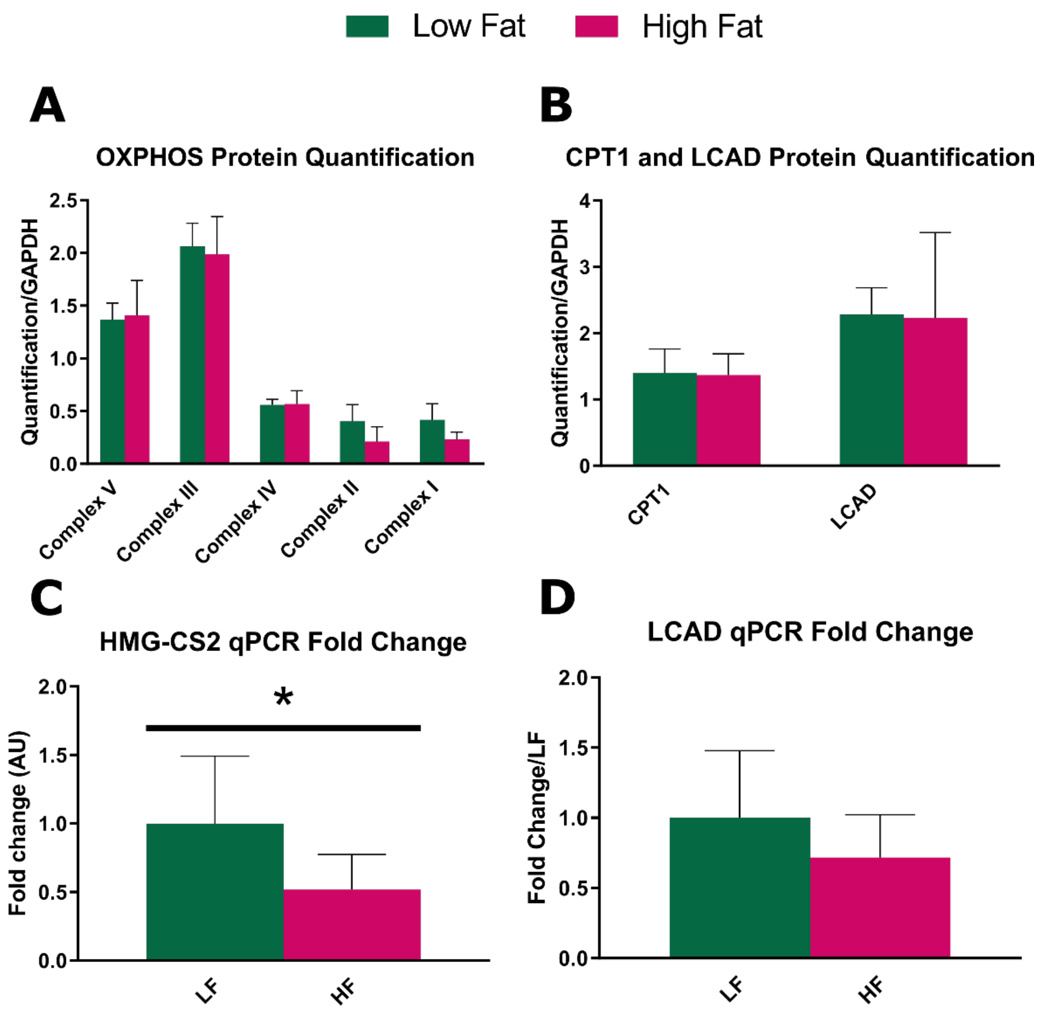


**Figure S4. Effect of 16 weeks of HFD on C57BL/6J mouse expression profile of electron transport chain, ketogenesis and fatty acid transport genes.** A) Western blot quantification of mitochondrial electron transport chain complexes 1-5 protein derived from freeze-clamped, powdered liver after perfusion. B) Western blot quantification of Carnitine palmitoyl transferase-1 (CPT1) and long chain acyl-CoA dehydrogenase (LCAD) extracted from perfused, freeze-clamped, powdered liver. C) Quantitative RT-PCR data for *HMG-CS2*, encoding Hydroxymethylglutaryl-CoA synthase 2, and D) *LCAD* obtained from RNA extracted from freeze-clamped, powdered liver after perfusion. Statistical significance was determined by Student’s t-test (*, $p < 0.05$).

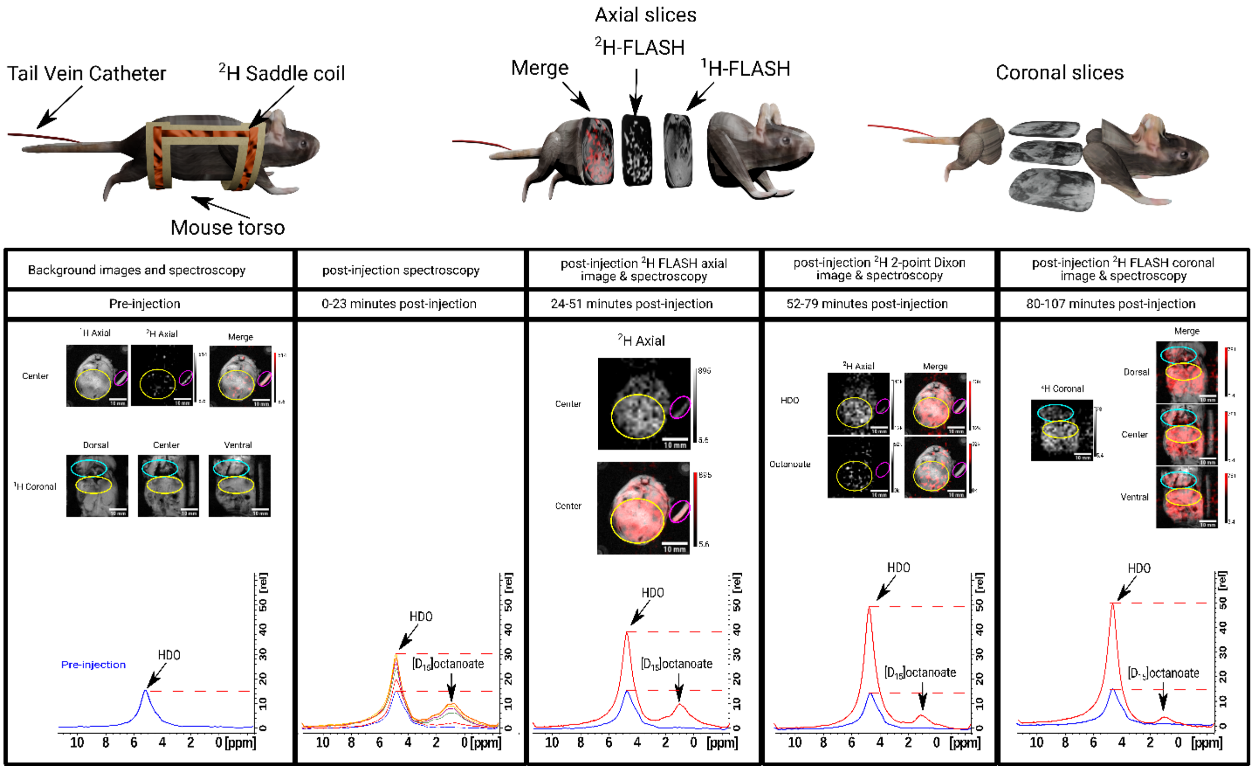


**Figure S5. Imaging timeline for axial and coronal deuterium MRI.** The top panel illustrates the physical location of the mouse (left) with the relative positioning of the deuterium saddle coil to the torso and the tail vein catheter. In the center the axial slices through the mouse torso are demonstrated by the overlay of the $^2$H axial slice and the $^1$H axial slice in order from left to right. Multiple coronal slices taken across the mouse torso were represented on the far right of the top panel. The Bottom panels from left to right demonstrate the series of background images, post injection spectral acquisition and subsequent axial and coronal deuterium images.

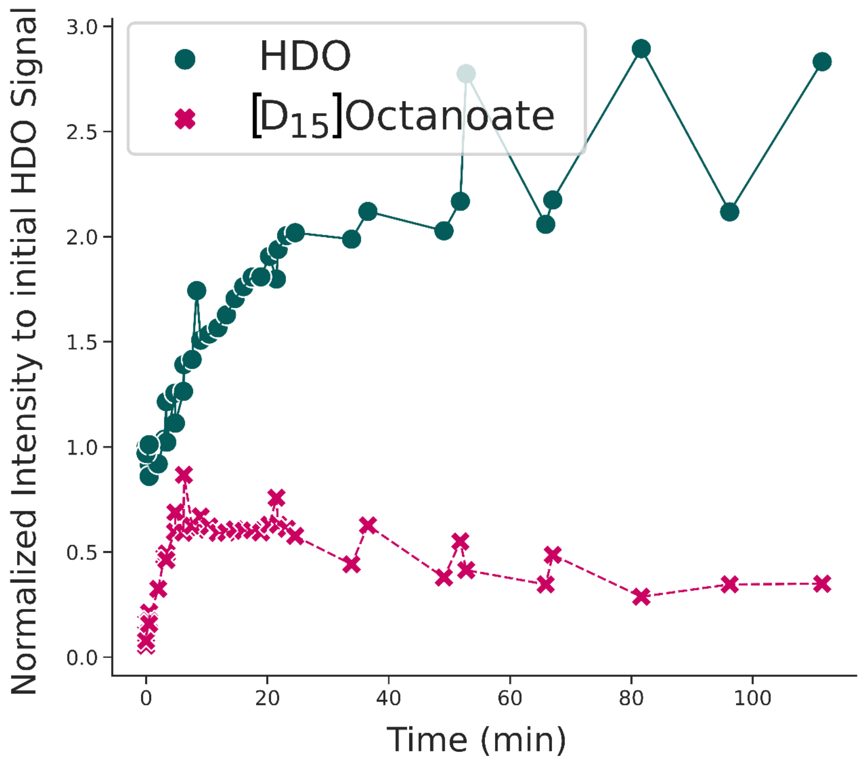


**Figure S6. Deuterium spectroscopy of mouse torso post [D$_{15}$]octanoate tail vein injection.** Quantified peaks from $^{2}$H spectra of three C57BL/6J mouse torsos post [D$_{15}$]octanoate injection (n=3) taken every 1.25 minutes for the first 25 minutes and between every image acquired subsequently.

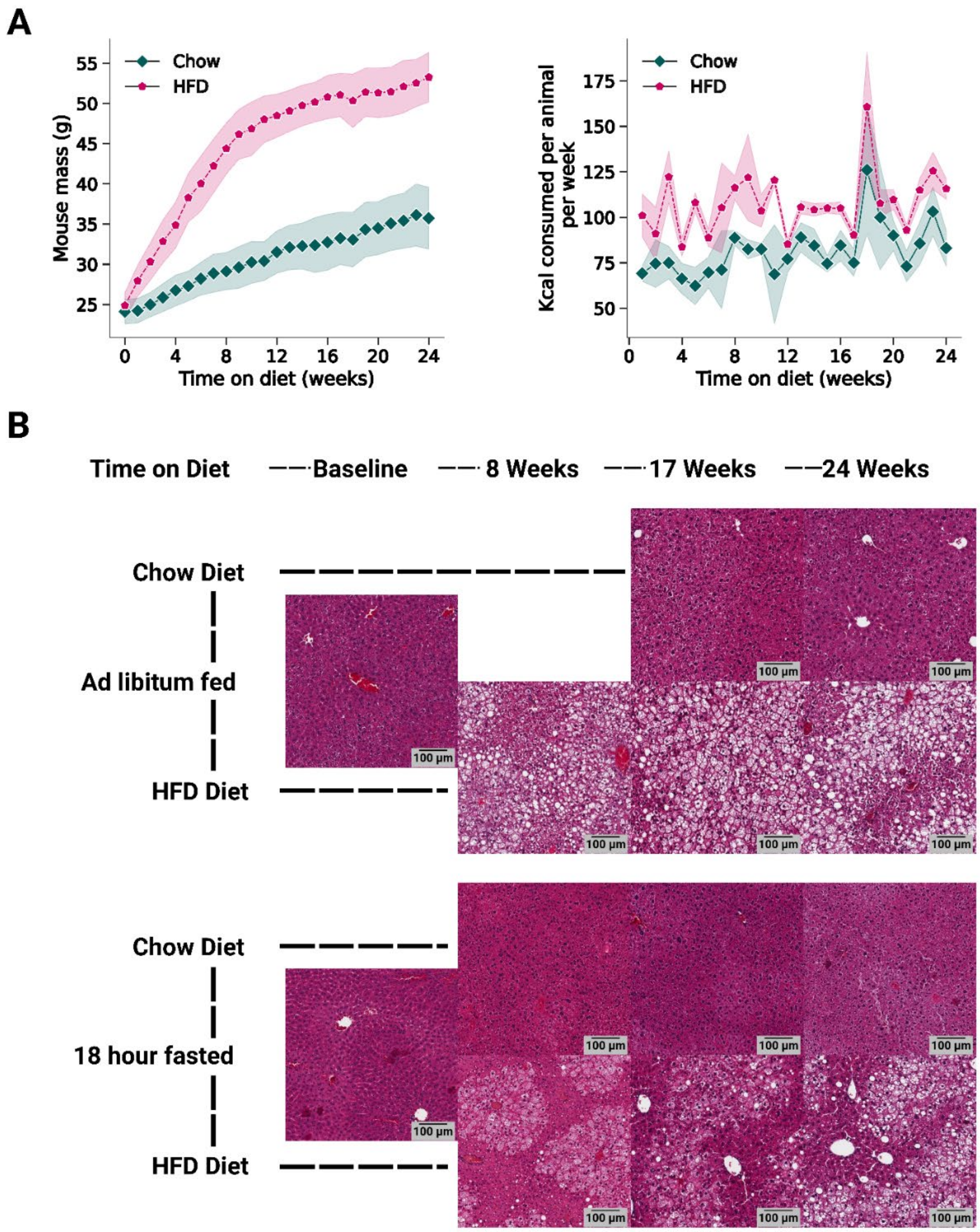


**Figure S7. Effect of HFD on body weight and liver.** (A) Graphs show mouse mass and calorie consumption for animals on the indicated diets that were used for DMRI imaging analysis in Figures 5 and 6. Diets were initiated with 8-week-old mice. Data are presented as average ± 95% confidence interval (shaded regions) for all mice on each diet. (B) Representative histological timeline of median lobe liver slices from baseline to 24 weeks of normal chow diet or HFD for *ad libitum* fed or 18-hour fasted mice. No image is shown for *ad libitum* fed chow diet at 8 weeks. Histology was used to calculate steatosis shown in Figure 5.

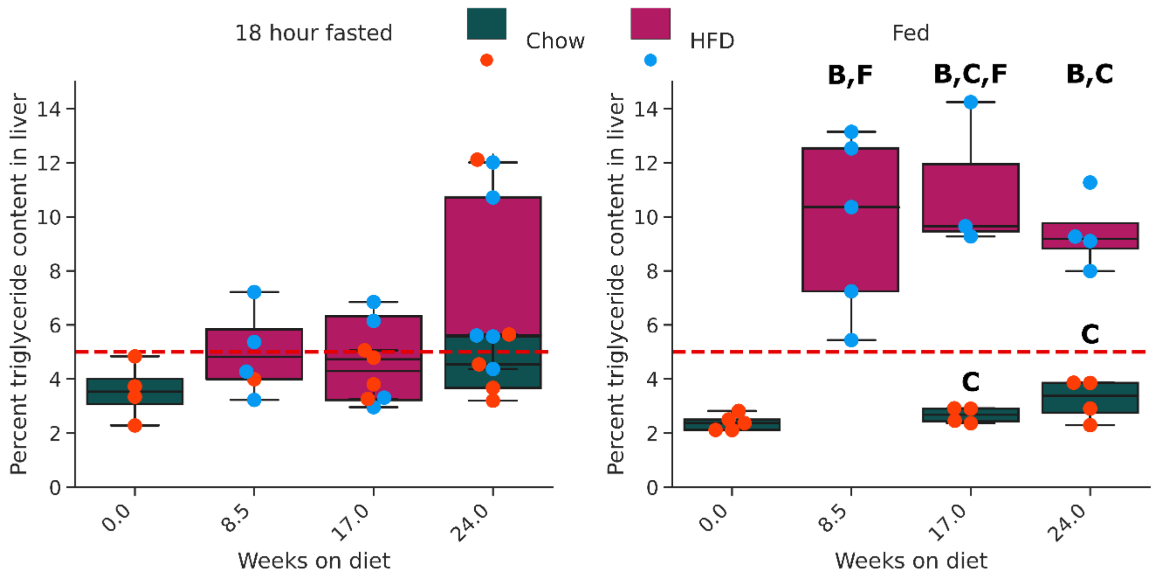


**Figure S8. Percentage triglyceride mass of the liver in C57BL/6J mice on HFD for 24 weeks. The red dashed line represents 5% hepatic triglyceride content by weight.** B refers to significance relative to baseline. C refers to significance when comparing the HFD mice to their chow diet counterparts for the same age. F refers to significance when comparing fed and fasted animals within the same diet and age.

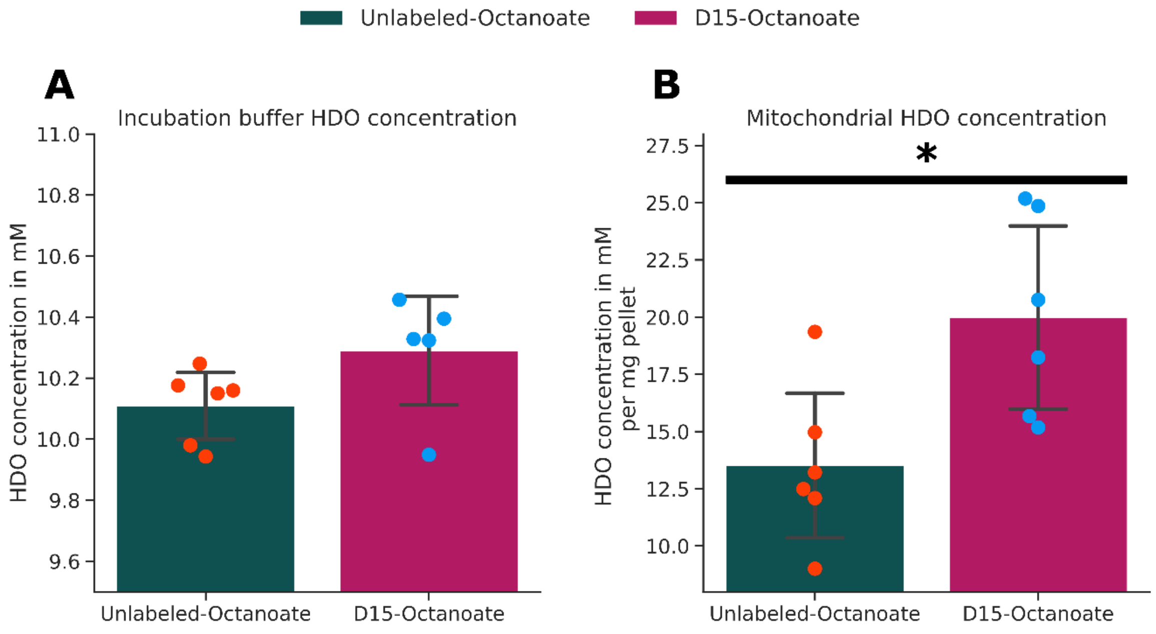


**Figure S9. HDO concentrations in mitochondria and buffer following incubation with [$D_{15}$]octanoate or unlabeled sodium octanoate.** A) The mitochondrial incubation buffer post incubation with [$D_{15}$]octanoate (dark pink) or unlabeled sodium octanoate (dark green). B) Mitochondrial pellets post 5-minute incubation with [$D_{15}$]octanoate (dark pink) or unlabeled sodium octanoate (dark green) and 1.5 hours of buffer washes. Data are presented relative to pellet mass. Statistical significance was determined by Student's t-test (*, $p < 0.05$).

**Table S1** Metabolic model used for INCA modeling with the atomic mapping for aliphatic deuterium and hydrogens and their transitions shown per reaction. For β-oxidation we emphasize that the deuterium mapping starts at the β position relative to the carboxyl carbon.

| |
|---|
| Pyruvate.perf (abc) -> Pyruvate (abc) |
| Lactate.perf (abcd) -> Lactate (abcd) |
| Lactate (abcd) -> Lactate.out (abcd) |
| Pyruvate (abc) -> Pyruvate.out (abc) |
| H (a) + H (b) <-> H2O (ab) |
| O2.in -> O2 |
| O2 -> O2.out |
| H2O (a) -> H2O.out (a) |
| **β-Oxidation** |
| Octanoate (abcdefghijklmno) + FAD + H (p) -> Octenoyl (cdefghijklmno) + FADH2 (pa) + H (b) |
| Octenoyl (abcdefghijklm) + H (o) -> Octanoyl (aobcdefghijklm) |
| Octanoyl (abcdefghijklmn) + NAD + H (o) -> NADH (c) + Hexanoate (defghijklmn) + AcCoa (abo) |
| Hexanoate (abcdefghijk) + FAD + H (l) -> Hexenoyl (cdefghijk) + FADH2 (al) + H (b) |
| Hexenoyl (abcdefghi) + H (j) -> Hexanoyl (ajbcdefghi) |
| Hexanoyl (abcdefghij) + NAD + H (k) -> NADH (c) + Butanoate (defghij) + AcCoa (abk) |
| Butanoate (abcdefg) + FAD + H (h) -> Butenoyl (cdefg) + FADH2 (ah) + H (b) |
| Butenoyl (abcde) + H (f) -> Butanoyl (afbcde) |
| Butanoyl (abcdef) + NAD + H (g) -> NADH (c) + AcCoa (abg) + AcCoa.t (def) |
| **Gluconeogenesis** |
| Pyruvate (abc) + NADH (d) <-> NAD + Lactate (abcd) |
| Pyruvate (abc) -> OAA (ab) + H (c) |
| OAA (ab) -> PEP (ab) |
| PEP (ab) + PEP (cd) + NADH (e) + NADH (f) -> glucose (abcdef) + O2 + NAD + NAD |
| **Electron Transport Chain** |
| NADH (a) + 0.5*O2 -> NAD + H (a) |
| FADH2 (ab) + 0.5*O2 -> FAD + H (a) + H (b) |
| **TCA Cycle** |
| Pyruvate (abc) + NAD + H (d) -> AcCoa (abc) + NADH (d) |
| AcCoa (abc) + OAA (de) -> Citrate (abde) + H (c) |
| Citrate (abcd) + H (e) + H (f) + NAD -> akg (abef) + NADH (c) + H (d) |
| akg (abcd) + NAD + H (e) -> Succinate (abcd) + NADH (e) |
| akg (abcd) <-> glutamate (abcd) |
| Succinate (abcd) + FAD + H (e) <-> Fumarate (ad) + H (b) + FADH2 (ce) |
| Fumarate (ab) + H (c) <-> Malate (abc) |
| Malate (abc) + NAD <-> OAA (bc) + NADH (a) |
| **Ketogenesis** |
| AcCoa.t (abc) + OAA (de) -> Citrate (abde) + H (c) |

| |
|---|
| AcCoa.t (abc) + AcCoa (def) <-> AcAc_coa (abcde) + H (f) |
| AcCoa (abc) + AcAc_coa (defgh) <-> HMG (defabgh) + H (c) |
| AcCoa (abc) + AcCoa (def) <-> AcAc_coa (abcde) + H (f) |
| HMG (abcdefg) + H (h) <-> AcAc (abcde) + AcCoa (hfg) |
| AcAc (abcde) + NADH (f) -> BHB (abcdef) + NAD |

### Table S2. Hepatic steatosis comparison.

| histology steatosis p value sheet | | | | |
|---|---|---|---|---|
| steatosis comparison | diet | timepoint (weeks) | feeding state | p-value |
| fed vs fasted | chow | 0 | fed-fasted | 4.4E-02 |
| fed vs fasted | HFD | 8.5 | fed-fasted | 1.5E-01 |
| fed vs fasted | HFD | 17 | fed-fasted | 3.2E-01 |
| fed vs fasted | chow | 17 | fed-fasted | 6.4E-02 |
| fed vs fasted | HFD | 24 | fed-fasted | 1.6E-01 |
| fed vs fasted | chow | 24 | fed-fasted | 5.9E-02 |
| diet comparison | HFD-Chow | 17 | Fasted | 8.7E-03 |
| diet comparison | HFD-Chow | 24 | Fasted | 1.0E-02 |
| diet comparison | HFD-Chow | 17 | Fed | 2.1E-05 |
| diet comparison | HFD-Chow | 24 | Fed | 1.1E-04 |
| baseline comparison | HFD | 0-8.5 | Fasted | 2.1E-02 |
| baseline comparison | HFD | 0-17 | Fasted | 8.4E-03 |
| baseline comparison | HFD | 0-24 | Fasted | 1.0E-02 |
| baseline comparison | Chow | 0-17 | Fasted | 2.9E-01 |
| baseline comparison | Chow | 0-24 | Fasted | 1.8E-01 |
| baseline comparison | HFD | 0-8.5 | Fed | 6.9E-02 |
| baseline comparison | HFD | 0-17 | Fed | 3.4E-05 |
| baseline comparison | HFD | 0-24 | Fed | 3.0E-04 |
| baseline comparison | Chow | 0-17 | Fed | 5.1E-01 |
| baseline comparison | Chow | 0-24 | Fed | 6.8E-02 |

### Table S3. Hepatic triglyceride comparison.

| triglyceride p value sheet | | | | |
|---|---|---|---|---|
| steatosis comparison | diet | timepoint (weeks) | feeding state | p-value |
| fed vs fasted | chow | 0 | fed-fasted | 1.1E-01 |
| fed vs fasted | HFD | 8.5 | fed-fasted | 3.2E-02 |
| fed vs fasted | HFD | 17 | fed-fasted | 3.6E-02 |
| fed vs fasted | chow | 17 | fed-fasted | 2.8E-02 |
| fed vs fasted | HFD | 24 | fed-fasted | 3.4E-01 |
| fed vs fasted | chow | 24 | fed-fasted | 1.9E-01 |
| diet comparison | HFD-Chow | 17 | Fasted | 6.1E-01 |
| diet comparison | HFD-Chow | 24 | Fasted | 4.4E-01 |
| diet comparison | HFD-Chow | 17 | Fed | 3.3E-02 |
| diet comparison | HFD-Chow | 24 | Fed | 6.7E-04 |
| baseline comparison | HFD | 0-8.5 | Fasted | 2.0E-01 |
| baseline comparison | HFD | 0-17 | Fasted | 3.1E-01 |
| baseline comparison | HFD | 0-24 | Fasted | 5.4E-02 |
| baseline comparison | Chow | 0-17 | Fasted | 3.5E-01 |
| baseline comparison | Chow | 0-24 | Fasted | 2.4E-01 |
| baseline comparison | HFD | 0-8.5 | Fed | 7.6E-03 |
| baseline comparison | HFD | 0-17 | Fed | 3.2E-02 |
| baseline comparison | HFD | 0-24 | Fed | 1.5E-03 |
| baseline comparison | Chow | 0-17 | Fed | 2.0E-01 |
| baseline comparison | Chow | 0-24 | Fed | 1.1E-01 |

**Table S4. Liver characteristics during NAFLD progression in response to HFD feeding**. The table includes HDO production/ mg [$D_{15}$]octanoate, HDO production/ mg [$D_{15}$]octanoate/g liver weight, liver mass (g), % steatosis, % triglyceride by mass. Data are presented as average ± SD.

| diet | fed state | Weeks on diet | HDO produced/mg [$D_{15}$]octanoate | HDO produced /mg [$D_{15}$]octanoate /g liver weight |
|---|---|---|---|---|
| Baseline | 18 hour fasted | 0 weeks | 310.16 ± 47.91 | 309.67 ± 47.00 |
| Chow | 18 hour fasted | 17 weeks | 314.17 ± 67.87 | 229.21 ± 11.50 |
| Chow | 18 hour fasted | 24 weeks | 300.52 ± 140.07 | 207.59 ± 78.36 |
| HFD | 18 hour fasted | 8.5 weeks | 208.75 ± 24.45 | 199.08 ± 6.29 |
| HFD | 18 hour fasted | 17 weeks | 507.99 ± 37.57 | 186.16 ± 14.59 |
| HFD | 18 hour fasted | 24 weeks | 413.21 ± 179.70 | 147.87 ± 37.57 |
| Baseline | *ad libitum* fed | 0 weeks | 232.20 ± 49.46 | 203.41 ± 28.34 |
| Chow | *ad libitum* fed | 17 weeks | 282.85 ± 42.00 | 185.91 ± 30.72 |
| Chow | *ad libitum* fed | 24 weeks | 317.44 ± 62.82 | 196.51 ± 41.69 |
| HFD | *ad libitum* fed | 8.5 weeks | 202.84 ± 22.45 | 134.29 ± 3.42 |
| HFD | *ad libitum* fed | 17 weeks | 430.51 ± 49.70 | 126.20 ± 22.10 |
| HFD | *ad libitum* fed | 24 weeks | 397.37 ± 36.86 | 106.86 ± 10.14 |

**Table S4 Continued.**

| diet | fed state | Weeks on diet | liver mass (g) | % steatosis | % triglyceride by mass |
|---|---|---|---|---|---|
| Baseline | 18 hour fasted | 0 weeks | 1.01 ± 0.05 | 0.91 ± 0.55 | 3.55 ± 1.06 |
| Chow | 18 hour fasted | 17 weeks | 1.36 ± 0.23 | 1.48 ± 0.82 | 4.23 ± 0.84 |
| Chow | 18 hour fasted | 24 weeks | 1.44 ± 0.28 | 2.56 ± 1.91 | 5.83 ± 3.63 |
| HFD | 18 hour fasted | 8.5 weeks | 1.05 ± 0.13 | 7.11 ± 2.87 | 5.02 ± 1.70 |
| HFD | 18 hour fasted | 17 weeks | 2.76 ± 0.58 | 20.80 ± 6.46 | 4.82 ± 1.97 |
| HFD | 18 hour fasted | 24 weeks | 2.72 ± 0.77 | 19.69 ± 6.48 | 7.66 ± 3.45 |
| Baseline | *ad libitum* fed | 0 weeks | 1.22 ± 0.10 | 3.13 ± 1.40 | 2.38 ± 0.30 |
| Chow | *ad libitum* fed | 17 weeks | 1.53 ± 0.11 | 3.93 ± 1.78 | 2.66 ± 0.29 |
| Chow | *ad libitum* fed | 24 weeks | 1.62 ± 0.04 | 5.44 ± 1.53 | 3.23 ± 0.77 |
| HFD | *ad libitum* fed | 8.5 weeks | 1.51 ± 0.13 | 14.00 ± 5.48 | 9.74 ± 3.33 |
| HFD | *ad libitum* fed | 17 weeks | 3.45 ± 0.41 | 24.78 ± 2.49 | 11.06 ± 2.77 |
| HFD | *ad libitum* fed | 24 weeks | 3.72 ± 0.18 | 25.93 ± 4.12 | 9.41 ± 1.36 |

**Table S5. Comparison of the effect of HFD and feeding state on hepatic β-oxidation measured by deuterium MRI.**

| Comparison | time on diet | diet | feeding state | p-value (total HDO) | p-value (relative HDO) | p-value liver mass |
|---|---|---|---|---|---|---|
| Relative to baseline | 0-8.5 | HFD | fasted | 5.94E-03 | 5.76E-03 | 5.36E-01 |
| Relative to baseline | 0-17 | HFD | fasted | 9.72E-03 | 2.73E-03 | 9.09E-03 |
| Relative to baseline | 0-24 | HFD | fasted | 3.38E-01 | 7.08E-04 | 2.09E-02 |
| Relative to baseline | 0-17 | Chow | fasted | 9.24E-01 | 1.65E-02 | 5.20E-02 |
| Relative to baseline | 0-24 | Chow | fasted | 9.02E-01 | 7.37E-02 | 5.29E-02 |
| Relative to baseline | 0-8.5 | HFD | fed | 3.48E-01 | 5.22E-02 | 4.08E-02 |
| Relative to baseline | 0-17 | HFD | fed | 1.31E-03 | 2.93E-02 | 1.05E-03 |
| Relative to baseline | 0-24 | HFD | fed | 1.97E-03 | 1.58E-02 | 7.71E-08 |
| Relative to baseline | 0-17 | Chow | fed | 1.71E-01 | 4.73E-01 | 5.64E-03 |
| Relative to baseline | 0-24 | Chow | fed | 5.69E-02 | 7.91E-01 | 1.76E-03 |
| Fed vs. Fasted | 0 | baseline | fed vs fasted | 5.16E-02 | 7.34E-03 | 1.50E-02 |
| Fed vs. Fasted | 8.5 | HFD | fed vs fasted | 7.55E-01 | 1.79E-05 | 8.62E-03 |
| Fed vs. Fasted | 17 | HFD | fed vs fasted | 1.69E-01 | 5.68E-03 | 1.05E-01 |
| Fed vs. Fasted | 24 | HFD | fed vs fasted | 8.73E-01 | 1.14E-01 | 7.94E-02 |
| Fed vs. Fasted | 17 | Chow | fed vs fasted | 4.68E-01 | 6.03E-02 | 2.68E-01 |
| Fed vs. Fasted | 24 | Chow | fed vs fasted | 8.34E-01 | 8.10E-01 | 2.82E-01 |
| HFD vs Chow | 17 | HFD vs Chow | fasted | 1.17E-02 | 4.08E-03 | 1.19E-02 |
| HFD vs Chow | 24 | HFD vs Chow | fasted | 3.63E-01 | 2.36E-01 | 3.79E-02 |
| HFD vs Chow | 17 | HFD vs Chow | fed | 4.22E-03 | 2.24E-02 | 1.62E-03 |
| HFD vs Chow | 24 | HFD vs Chow | fed | 4.67E-02 | 7.24E-03 | 4.61E-06 |

**Table S6. Metabolites derivatized by methoxyamine hydrochloride and dimethyl tert butyl silyl trifluoroacetamide (MBSTFA) and their associated quantitation ions for GC-MS identification.**

| Metabolite | m/z Quantitation Ion |
|---|---|
| Pyruvate | 174-179 |
| Lactate | 261-267 |
| Alanine | 260-264 |
| Glycine | 246-250 |
| 3-hydroxybutyrate | 275-279 |
| Succinate | 289-293 |
| Fumarate | 287-290 |
| a-ketoglutarate | 346-349 |
| Malate | 419-423 |
| Aspartate | 418-422 |
| Glutamate | 432-436 |
| Citrate | 459-463 |
